# Reconstructing prevalence dynamics of wildlife pathogens from pooled and individual samples

**DOI:** 10.1101/2023.11.02.565200

**Authors:** Benny Borremans, Caylee A. Falvo, Daniel E. Crowley, Andrew Hoegh, James O. Lloyd-Smith, Alison J. Peel, Olivier Restif, Manuel Ruiz-Aravena, Raina K. Plowright

## Abstract

Pathogen transmission studies require sample collection over extended periods, which can be challenging and costly, especially in the case of wildlife. A useful strategy can be to collect pooled samples, but this presents challenges when the goal is to estimate prevalence. This is because pooling can introduce a dilution effect where pathogen concentration is lowered by the inclusion of negative or lower-concentration samples, while at the same time a pooled sample can test positive even when some of the contributing samples are negative. If these biases are taken into account, the concentration of a pooled sample can be leveraged to infer the most likely proportion of positive individuals, and thus improve overall prevalence reconstruction, but few methods exist that account for the sample mixing process.

We present a Bayesian multilevel model that estimates prevalence dynamics over time using pooled and individual samples in a wildlife setting. The model explicitly accounts for the complete mixing process that determines pooled sample concentration, thus enabling accurate prevalence estimation even from pooled samples only. As it is challenging to link individual-level metrics such as age, sex, or immune markers to infection status when using pooled samples, the model also allows the incorporation of individual-level samples. Crucially, when individual samples can test false negative, a potentially strong bias is introduced that results in incorrect estimates of regression coefficients. The model, however, can account for this by leveraging the combination of pooled and individual samples. Last, the model enables estimation of extrinsic environmental effects on prevalence dynamics.

Using a simulated dataset inspired by virus transmission in flying foxes, we show that the model is able to accurately estimate prevalence dynamics, false negative rate, and covariate effects. We test model performance for a range of realistic sampling scenarios and find that while it is generally robust, there are a number of factors that should be considered in order to maximize performance.

The model presents an important advance in the use of pooled samples for estimating prevalence dynamics in a wildlife setting, can be used with any biomarker of infection (Ct values, antibody levels, other infection biomarkers) and can be applied to a wide range of host-pathogen systems.

## Introduction

When monitoring and studying pathogen transmission over time, it is essential to estimate prevalence dynamics. Prevalence, defined as the proportion of individuals in a population that tests positive for the current (e.g., presence of a pathogen or its genetic material) or past (e.g., antibody presence) presence of an infectious organism, is a key metric, yet can be difficult to estimate. The reason for this is that it is almost never feasible to test every individual in population, which means prevalence needs to be estimated from a population subset. As a result, methods are needed to estimate prevalence from imperfect data due to constraints in the number and quality of samples.

Sampling will depend on constraints (logistical, technical, individual availability, monetary), and different sampling strategies can be used to maximize the number of individuals being sampled [1, 2]. One such strategy is to pool samples, either by combining samples collected from different individuals (which reduces resource investments in testing and collection; [3]), or by collecting samples that already consist of material from multiple individuals (e.g., monitoring of SARS-CoV-2 in sewage; [4]). In studies of wildlife disease this latter approach is relatively common, for example when collecting fecal droppings in a den or cage containing multiple animals [2], or when collecting water samples in a lake or in wastewater [5]. An important drawback of the latter approach to pooling is that the sample cannot be linked to individual-level data, except indirectly under certain controlled conditions; [6].

Individual samples provide the highest-resolution information, as they allow additional individual-level data to be collected, including body measurements, estimates of sex and age class, and a wide range of biomarkers such as antibodies, blood proteins or other infections. These additional data are highly valuable as they can be used to learn more about correlates and drivers of infection. Depending on the study system, however, there can be several challenges to collecting and interpreting individual samples. A first is that the collection and processing of individual samples can be costly — in terms of effort, time or monetary resources — which limits sample sizes and temporal/spatial resolution. It can also be difficult to capture and sample individuals, for example when dealing with species that are elusive or live in low-density populations. Another challenge can arise when individuals do not shed a pathogen continuously but intermittently because of fluctuating pathogen concentrations. For example, the rodent *Mastomys natalensis* is known to shed arenavirus in varying concentrations [7]. Intermittent shedding means that it is possible to collect a negative sample or a sample with an undetectable pathogen concentration even though the individual can be considered infectious, leading to false negative results with regards to determining whether or not an individual is infectious.

A powerful study approach is to optimize the trade-off between sampling cost and data resolution by collecting both pooled and individual-level samples. This is commonly done in bat pathogen studies, where samples are collected from individual bats using net captures — which enables the collection of high-quality samples and associated individual variables — as well as from multiple bats simultaneously using plastic sheets placed under roosts [8–10]. This approach is particularly useful when the goal is to estimate prevalence dynamics.

When estimating prevalence, the use of pooled samples presents two well-known challenges, both resulting from the fact that multiple individuals contribute to the same sample. The first challenge is that a pooled sample can test positive regardless of how many of the contributing individual are actually positive. As a result, the proportion of positive pooled samples can be biased upwards, leading to over-estimates of prevalence [10]. The second challenge is the opposite of the first, and is the fact that a pooled sample can test negative even when one or multiple contributing individuals are positive. This can arise when the sample is diluted by negative samples, causing the concentration of the positive sample(s) to lower and fall below a detection threshold (which is called the dilution effect in pooled/group/composite testing literature; [11]). Assay sensitivity will be an essential factor in how low the diluted concentration can be before it can no longer be detected. Several approaches have been suggested to deal with these two challenges [11, 12], the most recent of which presents a Bayesian mixture model approach that can account for both at the same time under certain conditions [13]. Most studies on the analysis of pooled samples focus on testing protocols for cost reduction, with the goal of eventually identifying the positive individuals [3, 14, 15]. Perhaps for this reason, few methods have been developed for explicitly using pooled samples to estimate prevalence in the population [10,12,16–18], and even fewer have attempted to use the actual concentration of the infectious agent (or another biomarker like antibody concentration) in the pooled sample to estimate how many of the contributing individuals are positive [12, 13, 19]. A particular challenge arises when the underlying distribution of test values does not follow a standard-family (e.g. Gaussian) distribution, even though this is the most common situation, especially for wildlife populations [20, 21]. Few methods exist that can incorporate such distributions, and to our knowledge none provide a method for numerically calculating the full probability distribution of test values, instead using approximation methods [13, 19]. Leveraging the information present in the concentration of the infectious agent in pooled samples instead of only using binary negative/positive information can lead to significant improvements in the estimation of prevalence, particularly in the case of disease surveillance in wildlife populations.

We present a multilevel Bayesian modeling approach to estimate infection prevalence simultaneously from both individual and pooled samples, explicitly using the concentration of the infectious agent in pooled samples and thereby accounting for the biological mixing process that generates pooled sample concentrations. The model presents two key advances: first, the ability to estimate the false negative rate ensures that the effect coefficients of infection covariates can be estimated correctly, as these can otherwise be strongly affected by the presence of false negative samples. The second is the introduction of an algorithm that enables the full numerical calculation of the probability density function of concentrations in pooled samples.

Model use and performance is presented using simulated data inspired by a bat-pathogen study system, but we highlight that this approach can be used for any situation in which prevalence fluctuations are estimated from pooled samples with a known (or estimated) number of contributing individuals, especially when combined with individual samples. To illustrate the broader relevance, and test how the model performs under different conditions, we included relevant scenarios that each resemble a realistic biological situation. The approach presented here is particularly useful when the goal is not to identify which specific individuals are positive but to determine prevalence in the population, because there is no need to re-test depooled samples. Examples include monitoring SARS-CoV-2 prevalence [12, 18], estimating prevalence in wastewater if the number of contributing individuals can be estimated [5], assessing pathogen prevalence in the animal production industry [22], or estimating pathogen prevalence in wildlife populations [23]. Note that while the example presented here focuses on infection prevalence, the model can also be applied to other biomarkers such as antibodies.

## Methods

The main goal of this study is to estimate the true, unknown, proportion of pathogen-positive individuals over time, from both pooled and individual samples. Each of these types of samples presents a challenge for estimating prevalence, but also an opportunity, as outlined in Table 1. Note that the focus is on ”naturally” pooled samples, where collection was not done directly from individuals, as opposed to ”technically” pooled samples that were pooled intentionally after collection from individuals.

**Table 1:**
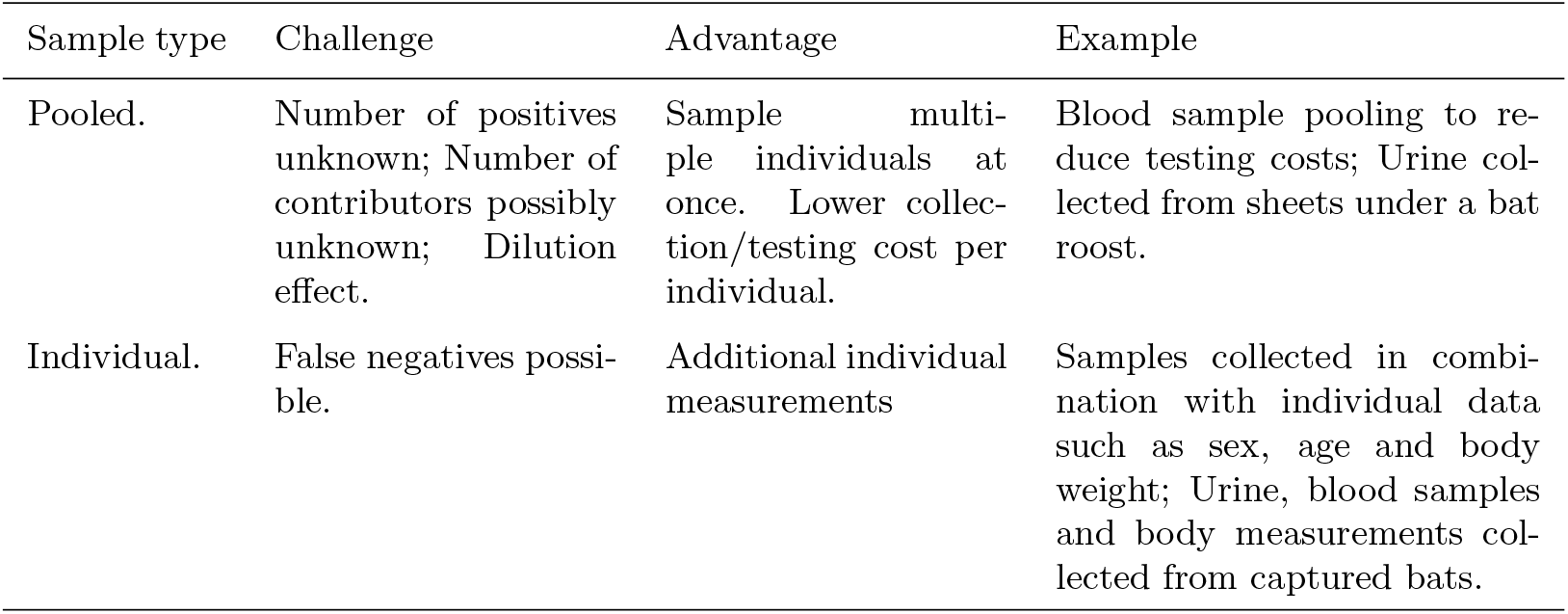
Sample types and their different challenges and advantages for estimating prevalence.

Here, we simulated data inspired by existing studies on flying foxes for research on temporal virus dynamics [8–10]. For the reasons mentioned above, bat virus studies often use field sampling designs that rely heavily on the collection of pooled urine and fecal samples under bat roosts [9]. A sampling design that incorporates pooled samples will be more beneficial for some wildlife species than for others, but there are no inherent limitations to which species this approach could be applied to. We chose to use simulated data only, as the goal of this study is to present and test a model to estimate prevalence, which can be done optimally when all underlying parameters are known and different scenarios can be generated. This makes it possible to determine how well the model is able to estimate the known parameters and prevalence dynamics for a range of scenarios. The simulated datasets are described below at the end of the Methods section.

The model is described in three parts, representing the multilevel/hierarchical nature of the model (1). The two main parts, a model for estimating prevalence from individual samples and a model for estimating prevalence from pooled samples, are linked by a third model of true, unobserved prevalence dynamics. We used a Bayesian multilevel model (also called a hierarchical model), as this provides a solid framework for linking the different model components, modeling unobserved latent parameters, incorporating prior knowledge through prior distributions, and providing posterior distributions of parameter estimates that show the uncertainty. While not done here, it would be straightforward to include an additional observation model that takes into account observation/measurement errors.

### Modeling individual samples

Individual (*i*) test result (negative or positive for biomarker presence) was modeled as a binary variable *y*_*i*_ (0 = negative, 1 = positive) using a Bernoulli distribution:

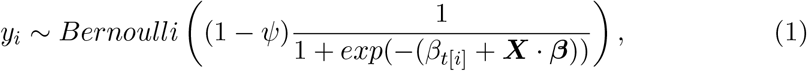

where *ψ* is the false negative rate that accounts for the lower prevalence resulting from the presence of false negative samples, and the remainder of the equation is a logistic regression, where *β*_*t*[*i*]_ is a varying intercept specific to each time at which individuals were sampled, ***X*** is a *n* x *k* matrix containing *k* covariates of *n* individuals, and ***β*** is a 1 x *k* matrix of regression coefficients. The logistic regression component allows estimating the correlation between individual-level covariates (e.g., biomarkers, age, body weight) and infection status. Prevalence 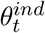 at each time point can in theory be calculated by taking the integral of *logit*^*−*1^(*β*_*t*[*i*]_ +***X*** *·****β***) over all covariates, but because this becomes highly computationally expensive when there is more than one covariate it is much more efficient to use a numerical approximation. Here, we used Monte Carlo integration [24], where *z* random samples are generated for each covariate from their distribution and the mean of the logistic function calculated at all sample combinations is the prevalence estimate 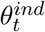 for time *t*:

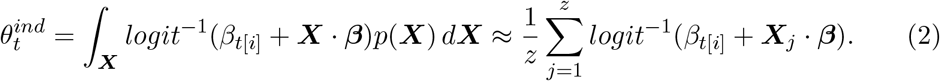

Random samples for each covariate are generated from their respective distributions. Here, covariates were modeled using a normal distribution, where the mean and standard deviation are included as parameters in the model. Finally, 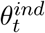 is used to estimate overall prevalence *θ*_*t*_ (as shown below in the section describing the true prevalence model).

The probability of an individual being positive, even when testing (false) negative, can be calculated using the inverse logit function,

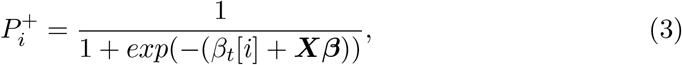

where individual shedding probability is informed by an individual’s covariate value(s) and prevalence at the time it was sampled. When there is a correlation between shedding status and one or more individual-level covariates, the predicted infection status can be used to identify which individuals may have tested false negative.

The prior distribution for *ψ* can be a beta distribution as it is bounded by 0 and 1. Because in many cases low false negative rates will be more likely, this could be a weakly informative distribution such as *Beta*(1, 2). The prior for *β*_*t*_ can be a weakly normal distribution such as *Normal*(0, 10). The prior distributions for the regression coefficients ***β*** will depend on the covariate and the way in which their correlation with shedding status is modeled, but in many cases this can be a weakly informative normal distribution such as *Normal*(0, 10) when using scaled covariates.

### Modeling pooled samples

The goal of this model is to estimate the proportion of positive bats using the Ct value of a pooled sample. The analysis of pooled samples can be challenging, leading to a large body of studies on pooled testing (also called group testing or composite testing, depending on the field) addressing the different problems related to pooled samples [11–13]. Most studies have focused on pooled testing in the context of laboratory assay cost reduction, where the main challenge is to find the optimal number of samples to pool given an expected proportion of positives [3]. An evolving challenge that is more applicable for understanding transmission dynamics is how to estimate the proportion of positive individuals. A number of approaches have been proposed for this, with many based on the model presented by [17]:

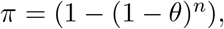

where *π* is the probability that a pooled sample tests positive, *θ* is prevalence, and *n* is the number of samples in the pool. Parameter *π* can then be used to model *Z ∼ Bernoulli*(*π*), where *Z* is a binary observed variable indicating whether or not the pooled sample is positive. This implementation has, for example, been proposed as a way to model prevalence dynamics over time for SARS-CoV-2, in combination with individual data [18]. This approach has two key limitations however. A first is that above a certain combination of pool size and prevalence (around 50%), most pooled samples will be positive, resulting in large uncertainty intervals surrounding the prevalence estimates. A second weakness is that this approach does not account for the fact that the concentration of pathogen is diluted by samples containing a lower concentration, including negative samples. This dilution effect has proven to be particularly difficult to address [19].

To date, most approaches have used binary test data for estimating prevalence using pooled samples [14, 16, 17]. Most assays, however, provide quantitative data, which are then turned into a binary negative/positive result based on a threshold value, and the additional information provided by the quantitative assay is lost. This quantitative information offers opportunities, however, that can address both limitations of the binary approach. Although few studies have developed methods to use the full quantitative test results for estimating prevalence from pooled samples [12, 19], the work by [13] in particular has shown how promising this approach can be. They used a Bayesian mixture model approach to estimate prevalence, taking into account the dilution effect based on the distribution of biomarker values (e.g. pathogen concentration) of negative and positive samples. A crucial part of these approaches is the use of a probability density function of positive test values. The methods in [12, 13] provide a useful approach for estimating these. To complement these approaches, we provide an algorithm to numerically calculate this probability density function so that it covers all possible combinations of numbers of positive and negative individuals while taking into account the underlying distribution of test values in the population.

We modeled pooled samples using their cycle threshold (Ct) value, a measure of the concentration of viral genetic material obtained using qRT-PCR (lower Ct value = higher concentration). The virus concentration in a pooled urine sample is determined by three key factors that influence the final pooled concentration: (1) proportion of positive bats, (2) concentration of virus shed by each positive bat, (3) relative urine volumes collected from each bat. Here we focus on the first two factors, and assume that the volumes collected from each bat are equal. In order to estimate the proportion of positive bats using the Ct value, it is necessary to calculate a probability distribution of Ct values for pooled samples, as this in turn enables calculating the likelihood of observing certain values given a combination of parameter values. A Ct probability distribution can be calculated by combining two key parts, a standard binomial probability density function (to take into account prevalence) and an ad-hoc distribution of probabilities of observing a pooled Ct value given a combination of negative and positive bats:

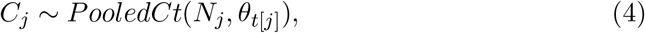

where *C*_*j*_ is the Ct value of pooled sample *j, N*_*j*_ is the total number of bats contributing to sample 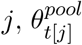 is prevalence at the time sample *j* was collected, and

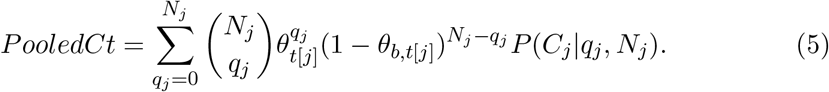

Here, 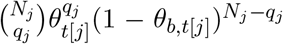 is the binomial probability of observing *q*_*j*_ positive out of *N*_*j*_ contributing individuals in pooled sample *j*, given a prevalence *θ*_*t*[*j*]_. *P* (*C*_*j*_|*q*_*j*_, *N*_*j*_) is the probability of observing Ct value *C*_*j*_ given *q*_*j*_ positive out of *N*_*j*_ individuals. *q*_*j*_ and *θ*_*t*[*j*]_ are the estimated parameters, while *N*_*j*_ and *C*_*j*_ are observed.

This equation closely matches equation 2 in [19].

Prior to model fitting, *P* (*C*_*j*_|*q, N*) must be calculated for each possible combination of *q, N*_*j*_ and *C*_*j*_, which is done according to the following algorithm:

1. Determine all possible combinations (with repetition) of *q* possible Ct values and *N*_*j*_ *− q* negative values.
2. For each combination:
  —2.1. Transform the Ct values of the positive samples to virus concentrations (conversion based on laboratory controlled testing, or testing of a range of individual samples).
  —2.2. Calculate the mean virus concentration.
  —2.3. Back-transform the mean virus concentration to its corresponding Ct value. Round up the Ct value to the next integer to mimic detection in RT-PCR (a concentration even slightly higher than a certain Ct value will not be detected until the next PCR cycle).
3. Count the number of combinations that result in Ct value *C*, and divide by the total number of combinations. This is Ct observation probability *P* (*C*_*j*_|*q, N*_*j*_), without accounting for prevalence in the population.

All code used for the calculation of the probability distributions can be found in Supplementary Information.

There are a number of important considerations when calculating *P* (*C*_*j*_|*q, N*_*j*_). A first is that while the algorithm assumes that each Ct value (in step 1) is equally likely, this is rarely the case. The distribution of Ct values in a population rarely follows a uniform distribution, and can instead follow many possible non-standard distributions (e.g., a skewed distribution when low concentrations are more likely). These distributions can also change over time and with changing biological conditions [25]. When this is the case, probability *P* (*C*_*j*_|*q, N*_*j*_) can be calculated by first calculating the total probability of each combination, then taking the sum of the total probabilities of all combinations that result in Ct value *C*, and dividing this by the sum of all total probabilities of all combinations. When the underlying Ct distribution changes over time, or under certain conditions, *P* (*C*_*j*_|*q, N*_*j*_) must be calculated for each of these situations. Individual samples, if collected, can be used to inform this distribution.

A second consideration is that urine volume is assumed to be equal for all *N* contributing bats. If this is not the case, the combinations can be corrected by normalizing for volume in the sample. This step requires knowledge of the volumes contributed by each individual. While this is possible in situations where samples are pooled after collection from individuals, this is unrealistic in field conditions. In this situation, the most parsimonious solution is to assume that all bats contributed equally to a pooled sample. This will of course rarely be the case, but variation in contributed volumes should not affect inference as long as it is not biased. Such biases could arise if infected bats, or bats shedding lower or higher virus concentrations, excrete different volumes than others. It is possible however to account for this when calculating *P* (*C*_*j*_|*q, N*_*j*_) if there is a model of how this bias occurs.

A third consideration is computational burden, which enforces a limit on the number of Ct values and the number of contributing bats. This is due to the fact that for each possible Ct value of a pooled sample, a probability is calculated for each possible combination 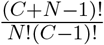 of Ct values *C* and individuals *N*. For example, in a simple situation where only 2 Ct values are possible, and a sample has 3 contributing individuals, the probability of observing a certain Ct value of the pooled sample must be calculated for 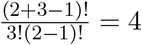 combinations. For more realistic numbers of 15 possible Ct values and 10 individuals, this becomes 1,961,256 combinations, increasing exponentially and rapidly reaching a maximum computationally feasible limit around combinations above 15 Ct values and 15 individuals. There are solutions for this, however. One solution would be to discretize Ct values into larger intervals (e.g., [21-24), [24-26), etc.), and/or setting all numbers of individuals above a certain maximum value equal to that value. This would lower the number of possible combinations and reduce computation time to feasible levels. Another solution, which would not require discretizing biomarker values or limiting the number of contributing individuals, would be to approximate the Ct probability distribution using Monte Carlo simulation/sampling [26] to generate a large number of random combinations of all values (versus numerically calculating every possible combination). While these solutions are likely to still result in good prevalence estimates, this will depend on the situation and should be tested with simulations prior to model fitting. We recommend taking these pool size requirements into account during the field experimental design process.

A full working example of the procedure to calculate the Ct probability distribution is provided in Figure 2.

**Figure 1:**
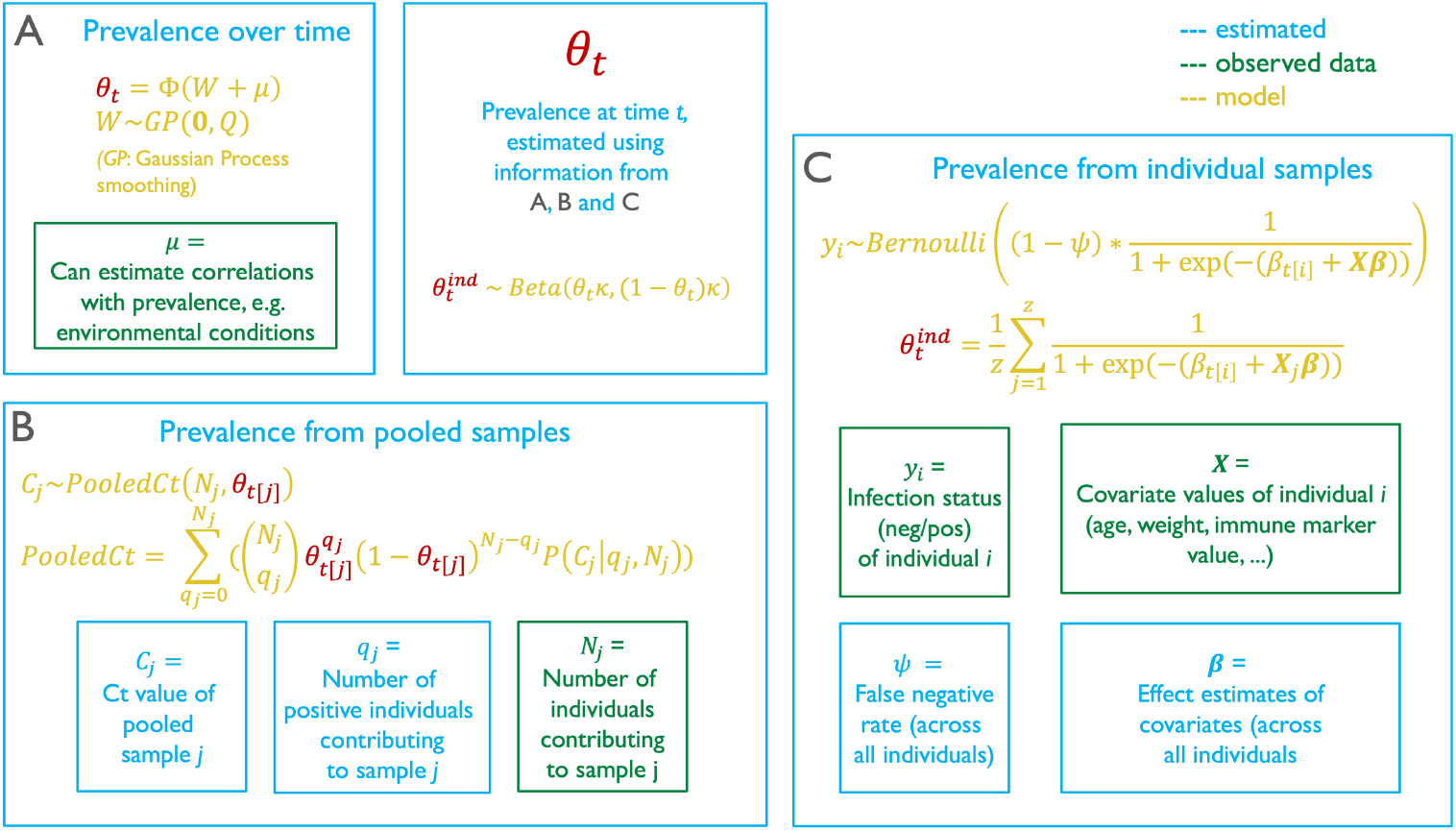
Multilevel model. A Gaussian Process model of prevalence (A) ensures that prevalence is estimated smoothly over time, using information about prevalence from the two other models (highlighted in red). Model (A) is able to test correlations between prevalence fluctuations and other variables such as temperature and precipitation. Model (B) illustrates the model that estimates prevalence from pooled samples, using the Ct value and number of contributing individuals as input data. Model (C) uses individual-level data to estimate prevalence, and enables estimating correlates of infection status.

**Figure 2:**
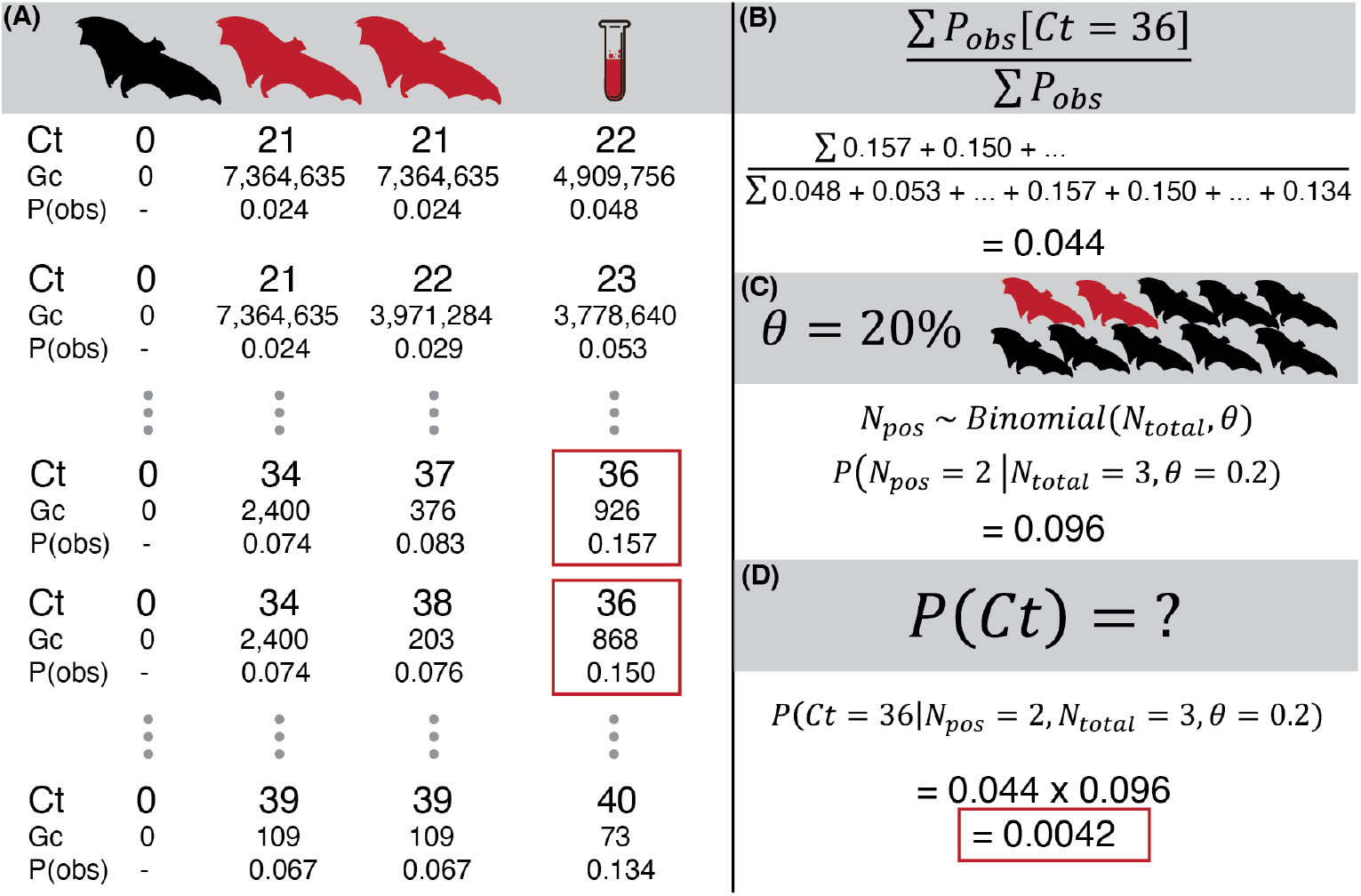
Illustration of how the probability of observing a Ct value in a pooled sample is calculated. In the example, we want to calculate the probability of observing a Ct of 36 with 2 out of 3 positive bats and 20% prevalence in the population. First (A), the pooled Ct value is calculated for every possible combination (with repetition) of 1 negative and 2 positive bats. For each combination, the Ct values (ln scale) are converted to genome copies (Gc) so that the pooled concentration can be calculated on a linear scale. The pooled genome copy concentration is then converted back to a Ct value, rounding up to the next integer to emulate the RT-PCR detection process. Next, for each combination of Ct values the corresponding probability of observing the pooled value is calculated by summing the respective individual probabilities that are estimated from the Ct distribution in individual bats. The probabilities corresponding with the target value of 36 are then summed and divided by the sum of all probabilities, to get an overall probability of observing Ct 36 (B). This probability is multiplied by overall prevalence in the population. The probability of observing 2 out of 3 positive individuals given a prevalence of 20% is then calculated (C) and multiplied by the probability of observing Ct 36 to get the final Ct probability given 20% prevalence and 2 out of 3 positive individuals (D). This example was randomly chosen for illustration purposes, and these steps are repeated for each possible combination of Ct values and contributing individuals.

### Modeling true prevalence in the population over time

The final model component is a model of *θ*_*t*_ dynamics, which explicitly incorporates the information about prevalence from individual and pooled samples through their respective models. This is possible because prevalence parameter *θ*_*t*_ is leverages information from both models, which further ensures that each of those two model components can benefit from the information about prevalence contained in the other. For pooled samples this is done as part of the Ct probability distribution, while for individual samples this is done using a hierarchical structure, where

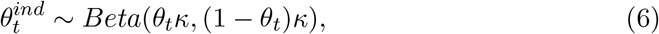

where *κ* is the precision parameter for the beta distribution, modeled using a weakly informative prior such as *Gamma*(2, 0.04). Prevalence *θ*_*t*_ changes over time *t* in a smooth way where prevalence will be more similar for times that are close together than for those that are farther apart. This temporal autocorrelation can be modeled in a variety of ways, and the choice of which model to use will depend on the research questions of interest. If the goal is to estimate prevalence dynamics over time, relatively simple smoothing functions can be used such as splines, weighted average or kernel functions [27]. If the goal is to model the underlying biological dynamics, it will be necessary to develop a more complex transmission model [28]. Here, we used a relatively simple Gaussian Process (GP) smoothing function, which uses a Gaussian kernel to model prevalence over time. This approach was based on the one used in [18].

A GP is a time continuous stochastic process *{X*_*t*_*}*_*t∈τ*_ where the set of variables *X*_*t*_ = (*X*_*t*1_, …, *X*_*tn*_)^*τ*^ is a multivariate Gaussian random variable (i.e., every combination of (*X*_*t*1_, …, *X*_*tn*_) has a univariate Gaussian distribution). Because *θ*_*t*_ *∈* [0, 1], a transformation must be used to map the real support of *X*_*t*_ to the [0, 1] interval, for which we used the inverse probit function Φ(*·*). We did this by modeling a latent prevalence process *W* := *{W*_*t*_*}*_*t∈τ*_ and transforming this to prevalence *θ*_*t*_ = Φ(*W*_*t*_). As prevalence and the form of the unobserved dynamic process are unknown, we used a GP prior on *W* with a covariance function that enables interpolation of prevalence over time (i.e., smoothing). There are multiple options for suitable covariance functions. Here, we used the exponentiated quadratic covariance function, which includes parameters for both the amplitude (lengthscale *ℓ*) and the oscillation speed (*σ*^2^) of the smoothing process,

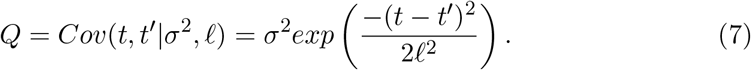

*W*_*t*_ thus becomes *W*_*t*_ *∼ GP* (**0**, *Q*), a zero-mean GP that allows independent modeling of the mean, which is useful for modeling the effect of covariates on prevalence, as *θ*_*t*_ becomes *θ*_*t*_ = Φ(*W*_*t*_ + *μ*), where *μ* can be any regression model.

A useful property of the covariance function is that by fitting the lengthscale parameter (ℓ), we can learn from the data how prevalence covaries over time is: the covariance between prevalence values separated by a time interval ℓ will be exactly 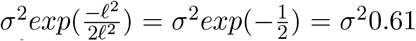, for an interval of 2ℓ this will be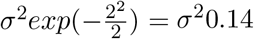, and so on.

The prior distributions for parameters *σ* and ℓ can be any continuous positive distribution. We used a truncated normal distribution for *σ* (*Normal*(0, 1), with 0 as lower bound for sampling) and an inverse gamma distribution for ℓ (*InverseGamma*(2.5, 150)). All priors used for model fitting can be found in the code in Supplementary Information.

### Testing model performance using simulated data

To test how well the model can estimate parameters under various circumstances, we simulated datasets that resemble realistic infection sampling scenarios. These datasets consisted of individual-level samples (collected directly from captured bats) and pooled urine samples (collected using a sheets under a roost), collected at certain time intervals (e.g., [8–10]). We created a main simulated dataset that resembles a common situation with regards to sample size and temporal resolution and was used as a point of reference for all analyses. To test model performance in different scenarios this main dataset was adapted in a number of ways that are described below.

For the main dataset, an autocorrelated fluctuating prevalence time series was generated for a time period of 300 (an arbitrary number, where the unit can be, but is not restricted to, days) time points using a b-spline function with knots at times 1, 100, 200 and 300. Coefficients for the b-spline function were chosen so that the function would result in reasonable prevalence fluctuations, not based on a specific system but useful for testing model performance under a range of sample availability scenarios (Figure 3). Ten sampling sessions were selected to occur evenly between times 1 and 300. At each sampling session, 50 individual-level catch samples and 50 pooled samples were generated. The infection status (negative/positive) of each individual sample was generated using a Bernoulli distribution with success probability equal to (1 *− ψ*)*logit*^*−*1^(*β*_*t*[*i*]_ + ***X*** *·* ***β***) (Figure 3A), where false negative rate *ψ* was set to 0.1 (i.e., 10% of positive samples test negative). The *β*_*t*_ values were generated by taking the logit of simulated prevalence at time *t*. One covariate was simulated by drawing random samples from a standard normal distribution. This covariate was then used to simulate outcome variables (infection status) for three different coefficients. Because the same *β*_*t*_ values were used for the three coefficients, this resulted in three different sets of outcome variables (Figure 3B), each with their own slightly different prevalence curve, which is a consequence of changes in prevalence due to the addition of *Xβ* to the intercept term.

**Figure 3:**
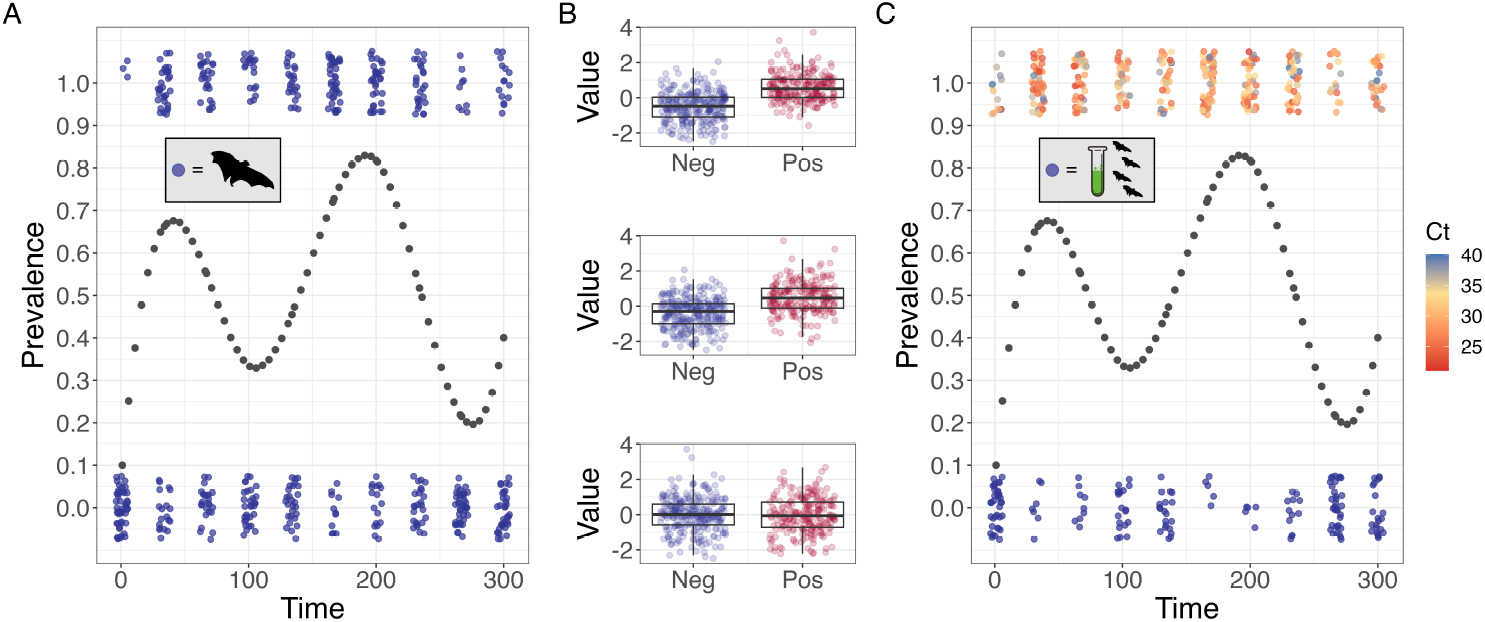
Simulated data. Black dots show true prevalence in the population, which was used to generate samples for 10 sessions over a period of 300 time points. Panel A shows individual negative (blue) and positive (red) samples, with false negative samples shown as red triangles. Panel B shows boxplots and data points for three simulated covariates for individual samples, with correlations being strong (top), moderate (middle) and random (bottom). Panel C shows pooled negative (blue) and positive (blue to red gradient corresponding with Ct value) samples. Note that infection data are binary (neg/pos) for individuals, and concentrations (Ct values) for pooled samples.

For each pooled sample, a Ct value was generated in four steps (Figure 3C). First, the number of bats contributing to the sample was simulated using a negative binomial distribution with size 30 and mean 2.3 (which results in a range between 1 and 10, with most numbers around 1 to 4). Next, each of the contributing bats was randomly assigned an infection status using a binomial distribution with success probability equal to prevalence at the corresponding sampling session. Then a Ct value was generated for each individual bat, with negative bats receiving a Ct value of 0 and positive bats receiving a Ct value randomly drawn from a non-standard, realistic probability distribution of Ct values. Last, the resulting Ct value of the pooled sample was calculated by first converting each individual Ct value to number of genome copies [25], calculating the mean number of genome copies (including the negative samples), and re-converting this mean of the pooled sample to a Ct value. Note that while a Ct value is generated for individuals contributing to a pooled sample, the individuals used for the ”individual sample” model described in the previous paragraph only have a negative or positive status, and not a Ct value. When required it is possible to add an observation process layer to the model that explicitly models the classification of sample into negatives or positives based on the concentration, as for example shown in [13, 19].

Main dataset simulation parameters are summarized in Table 2. Additionally, we show the importance of accounting for false negative individual samples when estimating covariate effects by fitting a model that does not include the false negative rate parameter. Last, to test model performance under different scenarios of data availability, we generated additional scenarios that are outlined in Table 3, including examples of when these scenarios can occur. Details and results for these scenarios are provided in Supplementary Information, including combined scenarios.

**Table 2:** Overview of the parameters used for the main simulated dataset.

| Parameter | Value |
| --- | --- |
| Times | 300 (arbitrary) time units |
| Number of sampling sessions | 10 |
| Timing of sampling session | Every 34 time units |
| Individual samples per session | 50 |
| Pooled samples per session | 50 |
| Infection data of individual samples | Binary (negative or positive) |
| Infection data of pooled samples | Concentration (Ct value) |
| False negative rate | 10% of positive individual samples tests negative |
| Individual covariate, strong correlation | Effect estimate = 3.3 |
| Individual covariate, moderate correlation | Effect estimate = 1.8 |
| Individual covariate, no correlation | Effect estimate = -0.06 |
| Ct distribution used to simulate pooled Ct values | Skewed low to high (details in Supplementary Information) |

**Table 3:** Simulated scenarios to test model performance. Details and model fit results are provided in Supplementary Information.

| Scenario | Details and examples |
| --- | --- |
| Pooled data only | Only pooled samples are used for model fitting. $\psi$ not estimated. This situation occurs when it is not possible not collect any individual data. Example: collecting wastewater samples for COVID-19 monitoring [4]. |
| Individual data only | Only individual samples are used for model fitting. $\psi$ not estimated. This situation is the most common when sampling populations. Example: cross-sectional sampling for monitoring arenavirus prevalence in rodents [29]. |
| Irregular sampling | The timing of sampling sessions is not regular, resulting in uneven time gaps between sessions. This situation occurs when regular sampling is not possible, or when sampling sessions need to be canceled due to conditions. Example: gaps in influenza A monitoring time series due to political instability [30]. |
| Low sample sizes | Lower sample sizes (20 instead of 50 for each sample type) per session. This situation occurs when it is not possible to sample many individuals. Example: logistically challenging captures of lions for canine distemper virus monitoring [31]. |
| Ct distribution mismatch | The distribution of Ct values used to calculate the likelihood for pooled sample Ct values is different from the true distribution used to simulate pooled Ct values. This situation can occur when the distribution of Ct values in the population is not well known. Example: small numbers of positive samples in individual bats make it difficult to describe the viral load distribution of filoviruses [32]. |
| Pool contribution count error | An error is added to the number of individuals contributing to a pooled sample. This situation occurs when it is difficult to count or estimate the number of individuals contributing to a pooled sample. Example: environmental sampling for *Leptospira* sp. prevalence estimation [33]. |
| Prevalence dynamics shape | A number of different, uncommon prevalence fluctuations are used to simulate the data. This situation occurs because prevalence dynamics can vary strongly depending on many factors. Example: measles prevalence dynamics exhibiting multi-annual cycles of varying magnitude [34]. |
| Prevalence covariate | A covariate that correlates with prevalence is estimated using Gaussian Process regression. Example: climate can drive inter-annual cycles of cholera transmission [35]. |

### Model implementation and code

All coding was done in R [36]. Model fitting was done with Stan [37] using R package rstan [38]. Plotting was done using packages ggplot2 [39], ggridges [40], patchwork [41] and Rcolorbrewer [42]. Prevalence splines were generated using the package splines [36]. Ct value probability distribution generation used the package Rccpalgos [43]. Supplementary information (including all code) is available online at https://doi.org/10.5281/zenodo.11520773.

## Results

Shedding prevalence dynamics estimated using the combined pooled and individual data closely matched the true dynamics, with true prevalence consistently falling within the posterior distribution (Figure 4A). All individual and prevalence covariate coefficients were estimated correctly except for the model excluding the false negative rate parameter, where the correct coefficient was 3.3 but the posterior mean estimate was 2.3 (95% CrI: 1.9-2.6). (Figure 4B-D). The model correctly estimated false negative rate (*ψ*) regardless of which covariate was used (Figure 4C).

**Figure 4:**
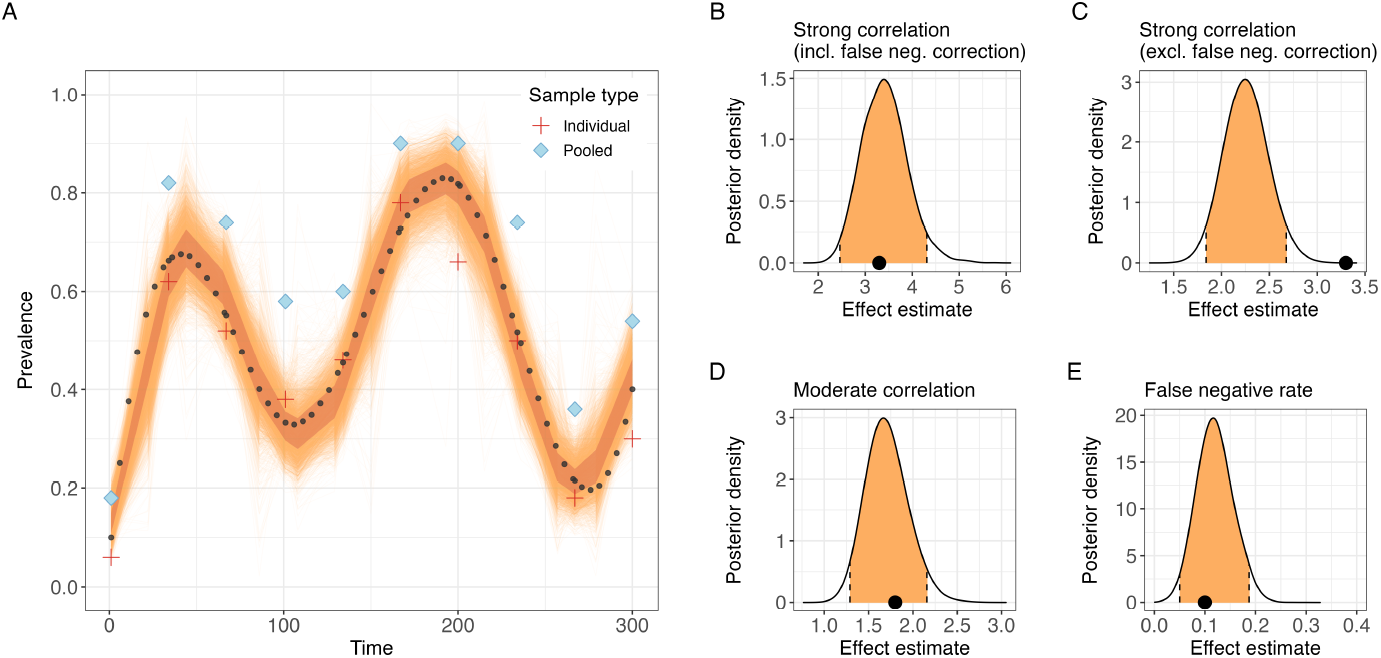
Model outputs for the main simulated dataset. (A) shows the distribution of fitted prevalence curves (cloud of 6,000 iterations from 5 chains) with 50% credible interval band overlaid. The black dots are the simulated prevalence values. The proportion of positive pooled and individual samples in each sampling session is shown using diamond and plus shapes, respectively. Panels (B) to (D) show the posterior distributions (95% credible intervals in orange) for two covariates (where B and C differ in whether or not false negatives were accounted for), and panel (E) shows the posterior distribution for the false negative rate *ψ*, with black dots indicating the true values.

When using only pooled data the model was still able to capture the true dynamics, while prevalence estimated using only individual data resulted in underestimates. (Figure 5A-B). The false negative rate could not be estimated in the absence of pooled data as there was no additional source of information to provide information about true prevalence over time.

**Figure 5:**
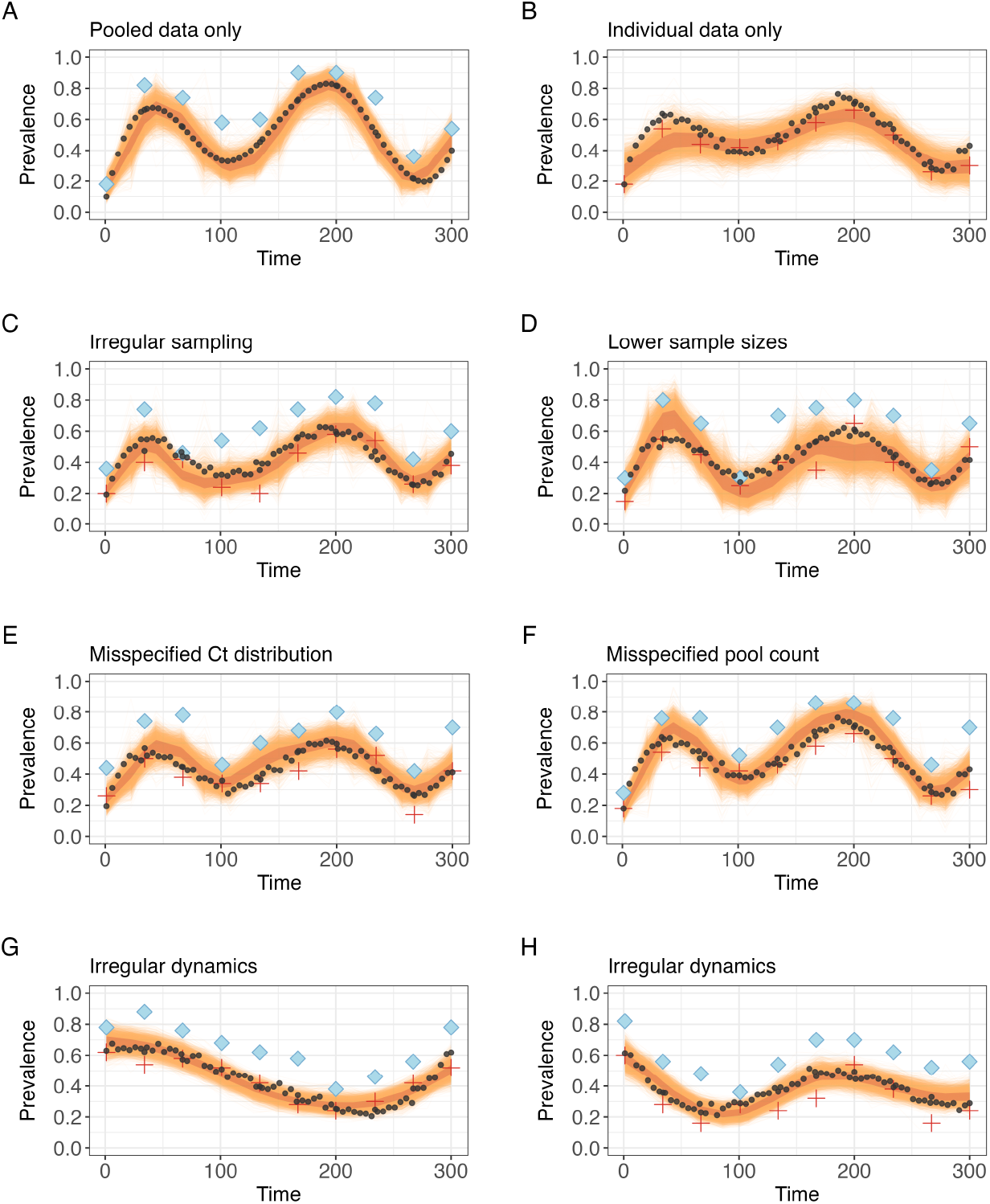
Fitted prevalence curves for different simulated scenarios. All scenarios used the same sample types, sizes and sessions as the main scenario shown in Figure 2, except where indicated. Session prevalence of pooled and individual data is shown using diamond and plus shapes, respectively. Black dots show true prevalence in the population. Specifics for each scenario are: (A) only pooled data; (B) only individual data; (C) sampling sessions are unevenly spaced over time; (D) lower sample sizes (20 per sample type) per session; (E) the Ct distribution used to simulate Ct values of pooled samples was not the same as that used to calculate the Ct probability distribution in the model, with the shapes inverted (i.e. low Ct values more likely); (F) an incorrect number of individuals contributing to a pooled sample was provided to the model for 50% of pooled samples; (G) and (H) data were simulated using irregular, unconventional prevalence dynamics.

When sampling sessions were timed irregularly, or when there were fewer sessions, prevalence was still estimated well but with a higher degree of uncertainty between larger time gaps (Figure 5C and Supplementary Information). For regular sampling with low sample sizes prevalence dynamics were still captured reasonably well overall, exhibiting increased variability that resulted in an over-estimation of the false negative rate parameter (Figure 5D). Asynchronous sampling of pooled and individual sessions resulted in prevalence dynamics that were very similar to those of the main simulated dataset (Supplementary Information). When combining irregular and asynchronous sampling with lower sample sizes or with fewer sampling sessions prevalence dynamics exhibited higher degrees of uncertainty due to the lower sample size or during larger time gaps without available samples (Supplementary Information).

A mismatch of the Ct distribution in the population (i.e., the Ct distribution used to construct *P* (*C*_*j*_|*q, N*_*j*_) did not correspond with the distribution used to simulate Ct values for pooled samples, see Supplementary Information for details) had a noticeable effect on the estimated prevalence dynamics (Figure 5E). Specifically, the model tended to overestimate prevalence, particularly during peaks, despite overall good performance. This effect was less pronounced when the distribution was less different from the true distribution (Supplementary Information). The model was not sensitive to moderately misspecified counts of the number of individuals contributing to a pooled sample (Figure 5F; 30% of the data were off by N = 1, 20% by N = 2), but was more strongly affected by large misspecifications (80% wrong by 1 or 2, 80% wrong by 1 to 5; Supplementary Information).

Last, the shape of the prevalence dynamics did not affect the model’s ability to estimate prevalence, as long as data were available to inform the fluctuations (5G and Supplementary Information). For example, the dynamics in Figure 5H have an initial peak that was not predicted by the model because this peak occurred between two sampling sessions.

## Discussion

Sample pooling offers major benefits through collecting data from multiple individuals at the same time, lowering costs for collection and testing, and enabling the use of samples that would otherwise be disregarded (such as sewage or fecal/urine under bat roosts or in animal dens) [4, 5, 9]. This study presents a Bayesian modeling approach that enables the estimation of prevalence dynamics from both pooled and individual samples by leveraging infection concentration of infectious agent in the pooled samples, allowing the distribution of infection concentrations to be any shape, and accounting for false negative results.

The model is able to successfully reconstruct prevalence dynamics for a wide range of eco-epidemiological scenarios. Model performance was tested for a range of relevant scenarios of infection dynamics and sampling schemes including irregular prevalence fluctuations, irregular timing of sampling and misspecified counts of individuals contributing to the pooled samples, and combinations of multiple scenarios. The model performs well when only one sample type was provided, which is particularly encouraging in the case of pooled samples, as it shows that field studies targeting only pooled samples would still allow precise reconstruction of prevalence dynamics. These results highlight the key strengths of the model: the explicit modeling of the mixing process in pooled samples allows accurate estimation of prevalence even when using only pooled samples, and the inclusion of pooled samples also enables correcting for the prevalence estimation bias in individual samples introduced by false negatives when both sample types are available. False negative rate is an epidemiological parameter commonly neglected in wildlife studies, yet important for inferring dynamics of infection at the individual level. In the model, estimation of false negative rates is made possible by the explicit integration of information about prevalence included in both data types. Importantly, accounting for false negative results ensures that covariate coefficients in individual-level regression models are estimated correctly, which we show would otherwise lead to estimation errors (Figure 4C). Lower sample sizes and large gaps between sampling sessions increased uncertainty, indicating that these are important factors to consider for study design.

The model introduces an algorithm to empirically calculate the probability distribution of observing a certain infection biomarker (here Ct from qRT-PCR) value given the estimated prevalence in the population and the number of individuals that contributed to the sample. This probability distribution enables the calculation of a likelihood for the biomarker values of the pooled samples. This approach for calculating a probability distribution can be adapted to other systems (e.g., analyzing pooled SARS-CoV-2 samples for monitoring prevalence) and other biomarkers (e.g., antibodies, blood chemistry). The approach can incorporate any non-standard family distribution of the biomarker. Encouragingly, we found that the model is quite robust against misspecifications of the underlying biomarker distribution. The calculation of this probability distribution function relies on a correct determination of the distribution of biomarker values in the population. We found that assuming a distribution that differs strongly from the real distribution can result in biased prevalence estimates. We therefore recommend an in depth prior exploration of biomarker distribution in the population, as well as a sensitivity analysis to assess how different realistic shapes of the distribution affect model output.

Prevalence reconstruction is a goal for many epidemiology and disease ecology studies, but this is often done as a necessary step towards learning what the drivers of pathogen transmission are. Such drivers can be intrinsic, such as individual immunity, herd immunity, individual variation in shedding, or behavior/movement (which can affect contact/transmission rates), or extrinsic, such as temperature and rainfall affecting pathogen survival, food availability affecting individual stress (which in turn affects immune competence, susceptibility and shedding). The modeling framework provides a way to incorporate and statistically test the effect of such covariates on the individual and the population/prevalence level. This enables testing of hypotheses about intrinsic or extrinsic drivers of infection, thereby contributing to a more mechanistic understanding of infection dynamics, beyond the phenomenological patterns. This also enables the development of models to predict prevalence.

The current model formulation has a number of requirements. Firstly, the model uses estimates of the number of individuals that contributed to a pooled sample. While the model is robust against moderately misspecified counts, we find that errors have to be within reasonable limits. However, when these counts are unknown or uncertain, this can be incorporated in the model by specifying a prior distribution of the number of individuals contributing to a pooled sample based on available data. A second model requirement is that the distribution of biomarker values, which are used to calculate the biomarker probability distribution of pooled samples, is assumed to be constant over time. Although this can be a reasonable baseline assumption, recent work suggests this may not always be the case [25]. Therefore, it is possible to adapt the model using a time-dependent probability distribution when pathogen shedding concentrations are known or suspected to be higher during certain periods. We recommend an in-depth analysis of the distribution of biomarker values in wild individual samples over time to determine whether the probability distribution used in the model needs to be time-dependent.

The model presented here provides a way to simultaneously leverage pooled and individual samples to accurately estimate the true underlying prevalence of infection in a population. It introduces a way to explicitly account for the biological mixing/dilution process in pooled samples, and ensures that individual covariate effects can be estimated correctly when false negative results are possible (this requires the use of both pooled and individual samples). The model is also shown to be robust against common issues associated with field-based data collection, such as observation noise and the often unknown shape of the underlying prevalence fluctuations. Crucially, this approach enables the accurate reconstruction of prevalence dynamics even when using pooled samples only, which is encouraging for designing lower-cost sampling strategies. The application of this model can directly enhance the efficacy and efficiency of bio-surveillance efforts by increasing inference and prediction. This is of particular interest in the case of wildlife that hosts pathogens of concern for human and animal health in geographical areas of high spillover risk.

## Acknowledgments

This work was funded by the U.S. Defense Advanced Research Projects Agency (DARPA) PREEMPT program Cooperative Agreement (D18AC00031), and the U.S. National Science Foundation (DEB-1716698; EF-2133763). The content of the information does not necessarily reflect the position or the policy of the U.S. government, and no official endorsement should be inferred. Thanks to Simon Brauer and Jacob Socolar for helpful feedback on the model.

