## Supplementary Information pdf for "Reconstructing prevalence dynamics of wildlife pathogens from pooled and individual samples" (from another version of this preprint: version 3, 2024-06-07)

[1]Wildlife Health Ecology Research Organization, San Diego, United States
[2]Department of Public and Ecosystem Health, College of Veterinary Medicine, Cornell University, Ithaca, United States
[3]Evolutionary Ecology Group, University of Antwerp, Antwerp, Belgium
[4]Department of Mathematical Sciences, Montana State University, Bozeman, USA
[5]Department of Ecology and Evolutionary Biology, University of California Los Angeles, Los Angeles, United States
[6]Centre for Planetary Health and Food Security, School of Environment and Science, Griffith University, Queensland, Australia
[7]Department of Veterinary Medicine, University of Cambridge, Cambridge, United Kingdom
[8]Department of Wildlife, Fisheries and Aquaculture, Mississippi State University, Mississippi State, United States
[*]

**Pooled data only** **14**

**Individual data only** **15**

**Irregular prevalence dynamics** **16**

**Lower sample size** **25**

**Fewer sampling sessions** **26**

**Misspecified Ct distribution** **28**

**Prevalence regression** **36**

**Asynchronous pooled and individual samples** **38**

**Irregular and asynchronous sampling times, lower sample sizes (20)** **39**

**Irregular and asynchronous sampling times, fewer sessions** **41**

**Additional figures** **43**

#### Model code

See Rmd file for Stan model code.

#### Ct probability distribution

Code used to generate Ct probability distribution $Ct_j \sim PooledCt(N_j, P_j)$:

```
# create the Ct probability array

# this is example code for creating the Ct probability
# array warning: long runtime

run.Ct.prob.again = F

if (run.Ct.prob.again) {

    library(RcppAlgos)

    # calculate Ct observation probabilities in pooled
    # samples, given a distribution of individual Ct value
    # probabilities

    range.Ntotal = 1:10  # range of possible total number of contributing individuals
    range.Npos = 0:10  # range of possible positive individuals
    range.Ct = c(0, 21:40)  # range of possible Ct values
    range.N.nonhosts = 0:10  # range of possible number of non-host individuals
    # amongst the contributing individuals

    # values of 0 in ct.prob.array were NA at first because
    # they don't exist, but stan complains when there are
    # NA's in arrays so need a value

    # function to calculate number of combinations
    n.comb.fun = function(n.elements, n.samples) factorial(n.samples +
        n.elements - 1)/(factorial(n.samples) * factorial(n.elements -
        1))

    # fast combination function used to calculate sums
    # below usage examples: comboGeneral(21:39,m=20,
    # repetition = T, lower = 1, upper = 10000)
    # comboGeneral(1:3,m=2, repetition = T, FUN =
    # function(x) mean(x), FUN.VALUE = 1.0) n.comb.fun(19,
    # 20)/1000 comboGeneral(21:39,m=20, repetition = T, FUN
    # = function(x) sum(x), FUN.VALUE = 1.0, lower = 1,
    # upper = 33578001) comboGeneral(21:39,m=10, repetition
    # = T, FUN = function(x) sum(x), FUN.VALUE = 1.0)

    # function to calculate number of permutations with
    # repetition, and multiply this by the sum of the
    # elements calculates the total probability of
    # observing a certain combination of Ct values, taking
    # into account the permutations example:
    # prod(factorial(as.numeric(table(c(1,1,1,2,3)))))
    n.perm.fun = function(x) factorial(length(x))/prod(factorial(as.numeric(table(x))))
    n.perm.sum.fun = function(x) sum(x) * (factorial(length(x))/prod(factorial(as.numeric(table(x)))))
```

```r
# ct combinations

# when running this function for the first time, this
# 'ct combinations' step needs to be done first it
# stores a list indicating which combinations of ct
# values can result in a given ct value

# for each number of positives calculate the sums of
# all possible ct (genome copy) combinations.  that
# list of sums will be used later instead of having to
# compute all combinations every time

for (i in range.Ntotal) {
    gc()

    # calculate sums of genome copy combinations
    cur.sums.gencop = comboGeneral(ct.to.gencop.fun(range.Ct[-1]),
        m = i, repetition = T, FUN = function(x) sum(x),
        FUN.VALUE = 1)

    saveRDS(cur.sums.gencop, paste0("genome.copy.sum.list.",
        i, ".RDS"))
    rm(cur.sums.gencop)
    print(i)
}

for (i in 1:length(range.Ct)) {

    gc()

    cur.ct = range.Ct[i]

    for (j in 1:length(range.Ntotal)) {

        cur.n.total = range.Ntotal[j]

        comb.possible.list = vector(mode = "list", length = cur.n.total)

        for (k in 2:(cur.n.total + 1)) {

            cur.n.pos = range.Npos[k]

            # read in sums of genome copies necessary
            # to import every time so that the memory
            # can be cleared (otherwise vector memory
            # exhausted at large npos), even though
            # this slows things down a bit
            cur.sums.perms = readRDS(paste0("genome.copy.sum.list.",
              cur.n.pos, ".RDS"))

            comb.possible.list[[(k - 1)]] = vector(mode = "list",
              length = length(range.N.nonhosts))
```

```
    for (l in 1:length(range.N.nonhosts)) {

      cur.n.nonhost = range.N.nonhosts[l]

      # calculate mean for each combination,
      # including the negatives and nonhosts
      # (i.e. divide by total number, not just
      # by number of positives)
      comb.means.gencop = cur.sums.perms/(cur.n.total +
        cur.n.nonhost)

      # convert means to ct values
      comb.means.ct = round(gencop.to.ct.fun(comb.means.gencop),
        1)  # rounding to 1 decimal digit because the conversion
      # can add some unexisting Ct val
      rm(comb.means.gencop)
      gc()

      # round up to nearest round Ct value
      # (reasoning = when doing PCR, a
      # concentration that is technically at
      # e.g. Ct 34.1 won't be detected at Ct =
      # 34, but will be detected at Ct = 35.
      comb.means.ct.round = ceiling(comb.means.ct)
      rm(comb.means.ct)
      gc()

      # transform all ct values larger than 40
      # to 0
      comb.means.ct.round[comb.means.ct.round > 40] = 0

      # determine combinations that result in
      # Ct values equal to that of the pooled
      # sample
      comb.possible.idx = which(comb.means.ct.round ==
        cur.ct)
      rm(comb.means.ct.round)
      gc()

      comb.possible.list[[(k - 1)]][[l]] = comb.possible.idx

      print(paste0("Ct = ", cur.ct, ", Npos = ",
        cur.n.pos, ", Ntotal = ", cur.n.total, " Nnonhost = ",
        cur.n.nonhost))
      rm(comb.possible.idx)
      gc()
    }
    rm(cur.sums.perms)
    gc()

  }
  saveRDS(comb.possible.list, paste0("comb.possible.list.Ct",
    cur.ct, ".Ntotal", cur.n.total, ".RDS"))
```

```
        rm(comb.possible.list)
        gc()
    }

}

# input Ct distribution in the population normally this
# would be created using empirical data, but here it is
# simulated low to high
ct.fit.prob = data.frame(ct = 21:39, prob = rev(dpois((21:39) +
    70, 24 + 70)/sum(dpois((21:39) + 70, 24 + 70))))
# ggplot(data = ct.fit.prob, aes(x = ct, y = prob)) +
# geom_line()

# flat: ct.fit.prob = data.frame(ct = 21:39, prob =
# rep(1/19,19)) ggplot(data = ct.fit.prob, aes(x = ct,
# y = prob)) + geom_line()

# high to low: ct.fit.prob = data.frame(ct = 21:39,
# prob =
# dpois((21:39)+70,24+70)/sum(dpois((21:39)+70,24+70)))
# ggplot(data = ct.fit.prob, aes(x = ct, y = prob)) +
# geom_line()

# center highest: ct.fit.prob = data.frame(ct = 21:39,
# prob = dnbinom(x = 21:39,size = 200,mu =
# 30)/sum(dnbinom(x = 21:39,size = 200,mu = 30)))
# ggplot(data = ct.fit.prob, aes(x = ct, y = prob)) +
# geom_line()

# run all code below for new individual Ct
# distributions

for (k in 2:length(range.Npos)) {

    cur.n.pos = range.Npos[k]

    if (cur.n.pos %in% 1:10) {
        # calculate sums of probabilities
        cur.sums.probs = comboGeneral(ct.fit.prob[, 2], m = cur.n.pos,
            repetition = T, FUN = n.perm.sum.fun, FUN.VALUE = 1)
    }

    saveRDS(cur.sums.probs, paste0("cur.sums.probs.npos",
        cur.n.pos, ".RDS"))
    print(cur.n.pos)
    rm(cur.sums.probs)
    gc()
}

ct.prob.array = array(data = 0, dim = c(length(range.Npos),
    length(range.Ntotal), length(range.Ct), length(range.N.nonhosts)),
    dimnames = list(Npos = range.Npos, Ntotal = range.Ntotal,
```

```r
        Ct = range.Ct, Nnonhost = range.N.nonhosts))

for (i in 1:length(range.Ct)) {

    cur.ct = range.Ct[i]

    for (j in 1:length(range.Ntotal)) {

        cur.n.total = range.Ntotal[j]

        comb.possible.list = readRDS(paste0("comb.possible.list.Ct",
            cur.ct, ".Ntotal", cur.n.total, ".RDS"))

        for (k in 1:(cur.n.total + 1)) {

            cur.n.pos = range.Npos[k]

            for (l in 1:length(range.N.nonhosts)) {

              cur.n.nonhost = range.N.nonhosts[l]

              if (cur.n.pos == 0 & cur.ct == 0) {
                ct.prob.array[k, j, i, l] = 1
              } else {

                if (cur.n.pos == 0)
                  cur.comb.possible.idx = integer(0)
                if (cur.n.pos > 0) {
                  cur.sums.probs = readRDS(paste0("cur.sums.probs.npos",
                    cur.n.pos, ".RDS"))
                  cur.comb.possible.idx = comb.possible.list[[cur.n.pos]][[l]]
                }
                if (length(cur.comb.possible.idx) == 0)
                  cur.prob = 0
                if (length(cur.comb.possible.idx) > 0) {

                  cur.prob = sum(cur.sums.probs[cur.comb.possible.idx])/sum(cur.sums.probs)

                }
                ct.prob.array[k, j, i, l] = cur.prob

              }
              print(paste0("Ct = ", cur.ct, ", Ntotal = ",
                cur.n.total, ", Npos = ", cur.n.pos, " Nnonhost = ",
                cur.n.nonhost))
            }

        }
    }

}

# low to high:
```

```
    saveRDS(ct.prob.array, "ct.prob.array.RDS")

} else {
    ct.prob.array = readRDS("ct.prob.array.RDS")
    ct.prob.array = ct.prob.array[, , , 1]
}
```

Ct frequency distribution used to calculate Ct observation probabilities:

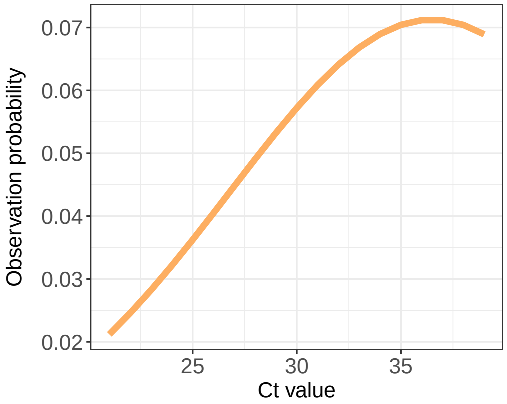

#### Main scenario

For the main simulated dataset, a fluctuating prevalence time series was generated for a time period of 300 days using a b-spline function with knots at days 1, 100, 200 and 300. Coefficients for the b-spline function were chosen so that the function would result in realistic prevalence fluctuations. Ten sampling sessions were selected to occur evenly between days 1 and 300. At each sampling session, 50 individual catch samples and 50 pooled samples were generated. The infection status (negative/positive) of each individual sample was generated using a Bernoulli distribution with success probability determined by prevalence at the session time and the covariate, including false negatives (10%).

For each pooled sample, a Ct value was generated in four steps. First, the number of bats contributing to the sample was simulated using a negative binomial distribution with size 30 and mean 2.3 (which results in a number of bats ranging between 1 and about 10, with most numbers around 1 to 4). Next, each of the contributing bats was randomly assigned an infection status using a binomial distribution with success probability equal to prevalence at the corresponding sampling session. Then a Ct value was generated for each bat, with negative bats receiving a Ct value of 0 and positive bats receiving a Ct value randomly drawn from a non-standard, realistic probability distribution of Ct values. Last, the resulting Ct value of the pooled sample was calculated by first converting the Ct value to number of genome copies, calculating the mean number of genome copies (including the negative samples), and re-converting to a Ct value.

#### Main model with strongly correlating individual covariate

##### Prevalence dynamics

Estimated prevalence dynamics for the main dataset, using a strongly correlating covariate. Black points = simulated prevalence. Orange lines = 5000 posterior draws. Orange band = 50% credible interval.

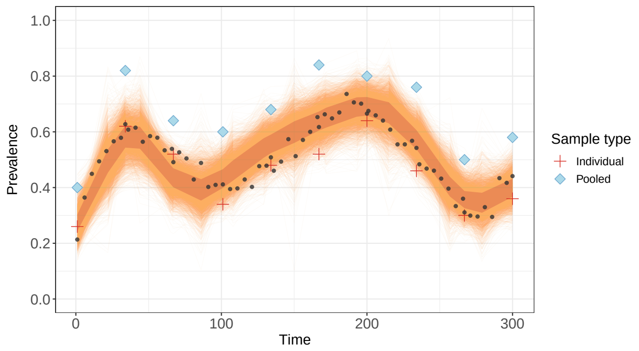

##### False negative rate

False negative rate posterior density with 95% CrI:
True value = 0.1.

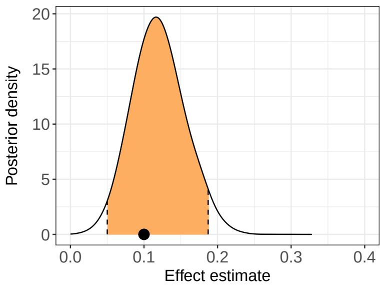

##### Individual covariate

Effect estimate posterior density with 95% CrI, strongly correlating covariate.
True coefficient = 3.3.

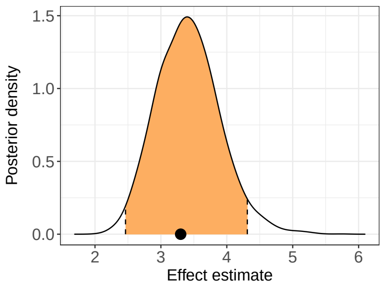

#### Main model excl. false neg rate, with strongly correlating individual covariate

##### Prevalence dynamics

Estimated prevalence dynamics for the main dataset, using a strongly correlating covariate. Black points = simulated prevalence. Orange lines = 5000 posterior draws. Orange band = 50% credible interval.

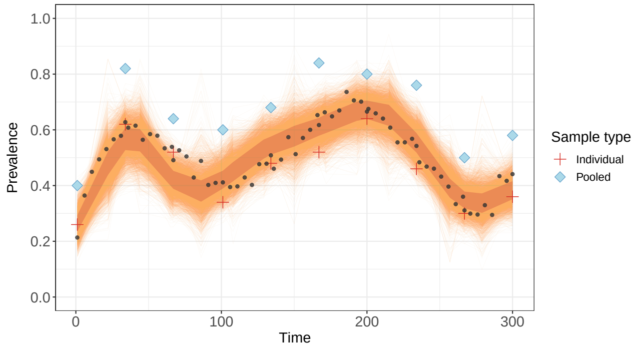

##### Individual covariate

Effect estimate posterior density with 95% CrI, strongly correlating covariate.
True coefficient = 3.3.

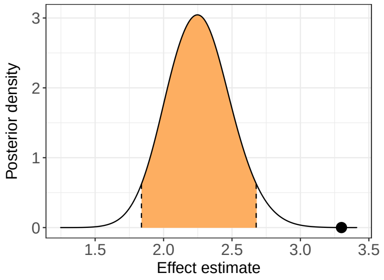

Posterior mean: 2.26 (95% CrI: 1.86-2.63).

#### Main model with moderately correlating individual covariate

##### Prevalence dynamics

Estimated prevalence dynamics for the main dataset, using a strongly correlating covariate. Black points = simulated prevalence. Orange lines = 5000 posterior draws. Orange band = 50% credible interval.

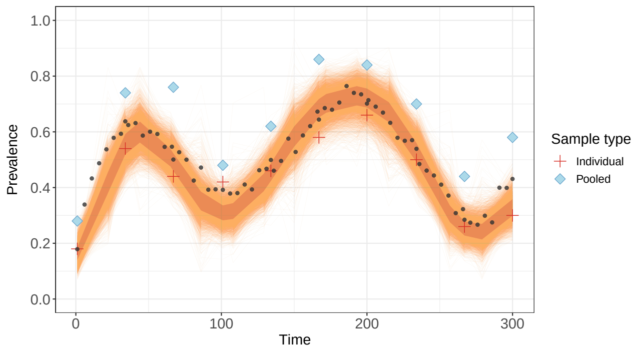

##### False negative rate

False negative rate posterior density with 95% CrI:
True value = 0.1.

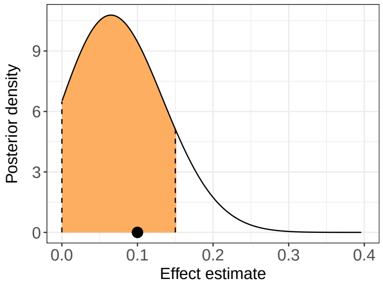

##### Individual covariate

Effect estimate posterior density with 95% CrI, strongly correlating covariate.
True coefficient = 1.8.

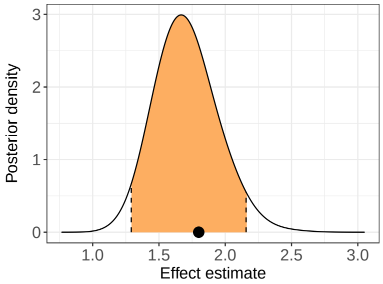

### Main model with non-correlating individual covariate

#### Prevalence dynamics

Estimated prevalence dynamics for the main dataset, using a strongly correlating covariate. Black points = simulated prevalence. Orange lines = 5000 posterior draws. Orange band = 50% credible interval.

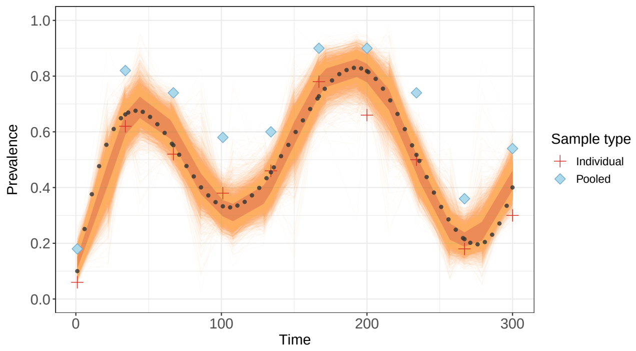

#### False negative rate

False negative rate posterior density with 95% CrI:
True value = 0.1.

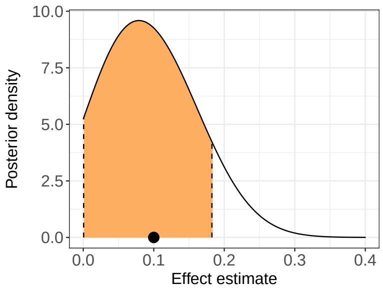

#### Individual covariate

Effect estimate posterior density with 95% CrI, strongly correlating covariate.
True coefficient = -0.06.

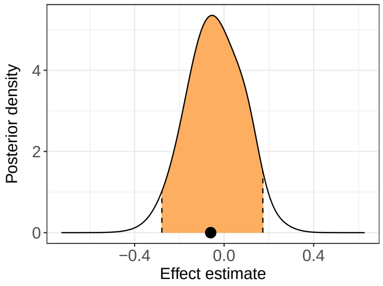

### Pooled data only

This simulated dataset is the same as the main dataset, but model fitting is done using pooled data only. This means that it is not possible to estimate individual shedding status or false negative rate.

#### Prevalence dynamics

Estimated prevalence dynamics for the main dataset, using pooled data only. Black points = simulated prevalence. Orange lines = 5000 posterior draws. Orange band = 50% credible interval.

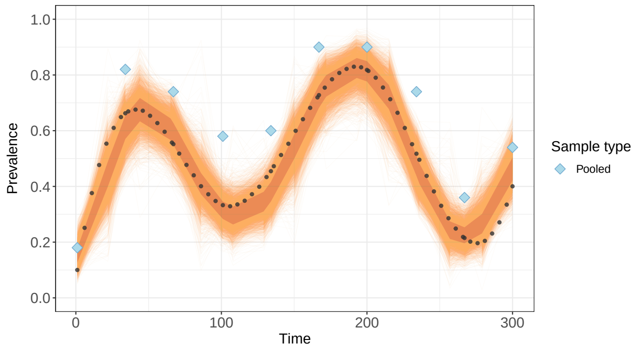

### Individual data only

This simulated dataset is the same as the main dataset, but model fitting is done using individual data only.

#### Prevalence dynamics

Estimated prevalence dynamics for the main dataset, using a moderately correlating covariate. Black points = simulated prevalence. Orange lines = 5000 posterior draws. Orange band = 50% credible interval.

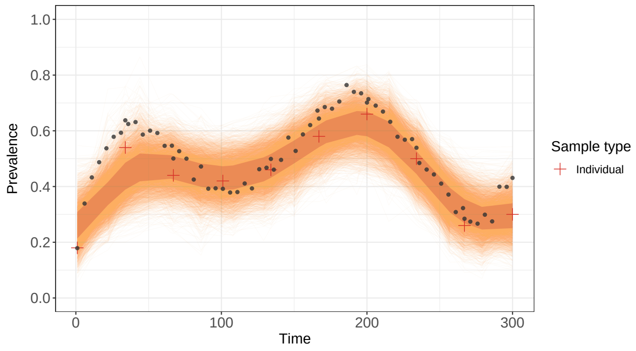

#### Individual covariate

Effect estimate posterior density with 95% CrI, moderately correlating covariate.
True coefficient = 1.8.

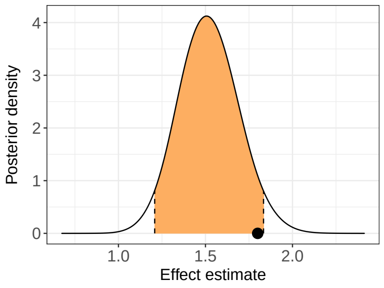

### Irregular prevalence dynamics

These datasets have the same sample structure (sizes and timing) as the main dataset, but prevalence dynamics were made more irregular.
The models use the weakly correlating individual covariate.
There are 5 versions of irregular dynamics, different each time, in order to assess model performance for different types of dynamics.

#### Irregular dynamics v1

##### Prevalence dynamics

Estimated prevalence dynamics for the main dataset, using a moderately correlating covariate. Black points = simulated prevalence. Orange lines = 5000 posterior draws. Orange band = 50% credible interval.

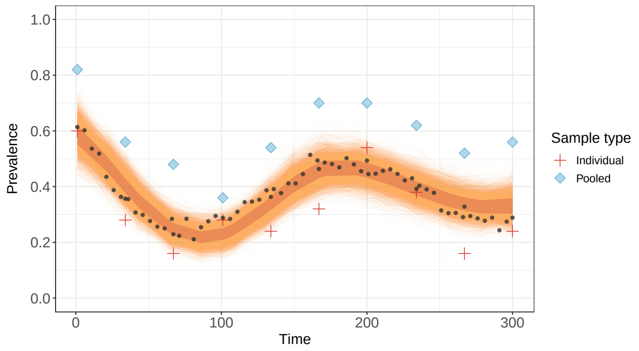

##### False negative rate

False negative rate posterior density with 95% CrI:
True value = 0.1.

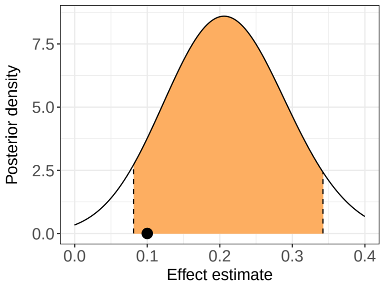

##### Individual covariate

Effect estimate posterior density with 95% CrI, moderately correlating covariate.
True coefficient = 1.6486208.

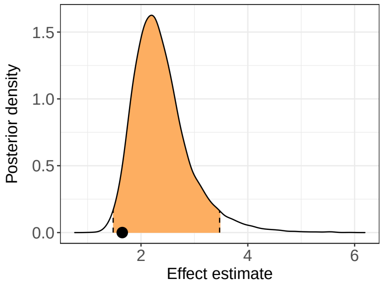

Posterior density
1.5
1.0
0.5
0.0
2
4
6
Effect estimate

#### Irregular dynamics v2

##### Prevalence dynamics

Estimated prevalence dynamics for the main dataset, using a moderately correlating covariate. Black points = simulated prevalence. Orange lines = 5000 posterior draws. Orange band = 50% credible interval.

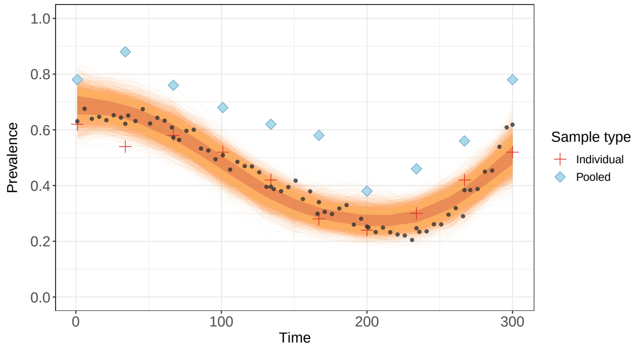

##### False negative rate

False negative rate posterior density with 95% CrI:
True value = 0.1.

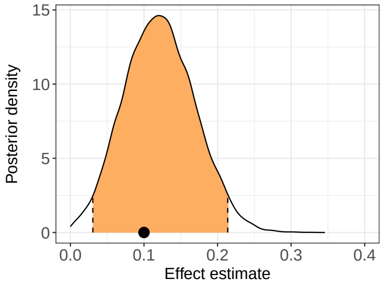

##### Individual covariate

Effect estimate posterior density with 95% CrI, moderately correlating covariate.
True coefficient = 2.0116604.

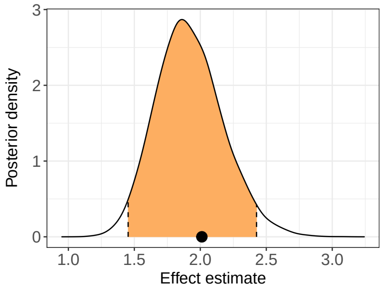

#### Irregular dynamics v3

##### Prevalence dynamics

Estimated prevalence dynamics for the main dataset, using a moderately correlating covariate. Black points = simulated prevalence. Orange lines = 5000 posterior draws. Orange band = 50% credible interval.

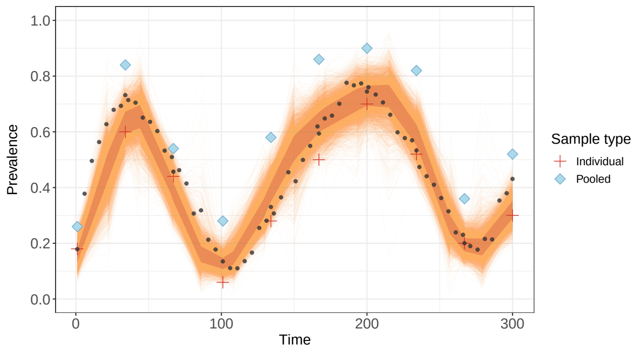

##### False negative rate

False negative rate posterior density with 95% CrI:
True value = 0.1.

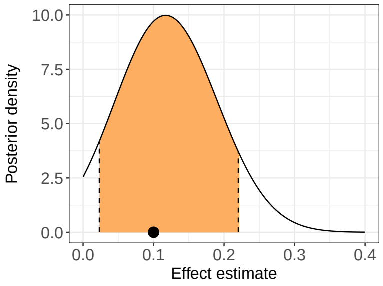

##### Individual covariate

Effect estimate posterior density with 95% CrI, moderately correlating covariate.
True coefficient = 1.9636848.

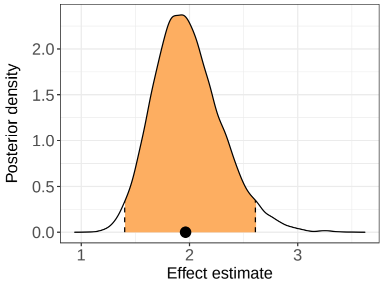

### Irregular dynamics v4

#### Prevalence dynamics

Estimated prevalence dynamics for the main dataset, using a moderately correlating covariate. Black points = simulated prevalence. Orange lines = 5000 posterior draws. Orange band = 50% credible interval.

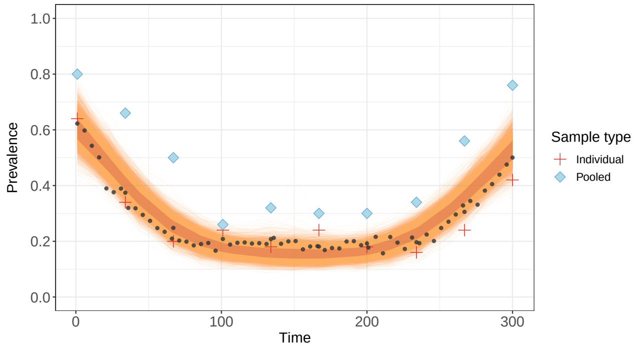

#### False negative rate

False negative rate posterior density with 95% CrI:
True value = 0.1.

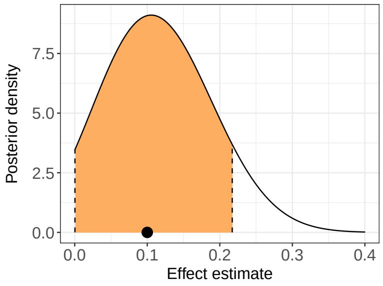

#### Individual covariate

Effect estimate posterior density with 95% CrI, moderately correlating covariate.
True coefficient = 2.0195733.

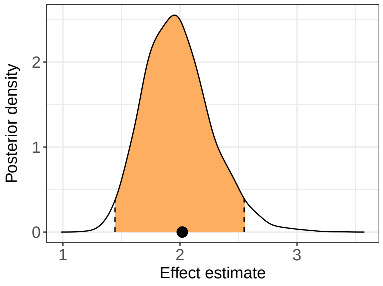

#### Irregular dynamics v5

##### Prevalence dynamics

Estimated prevalence dynamics for the main dataset, using a moderately correlating covariate. Black points = simulated prevalence. Orange lines = 5000 posterior draws. Orange band = 50% credible interval.

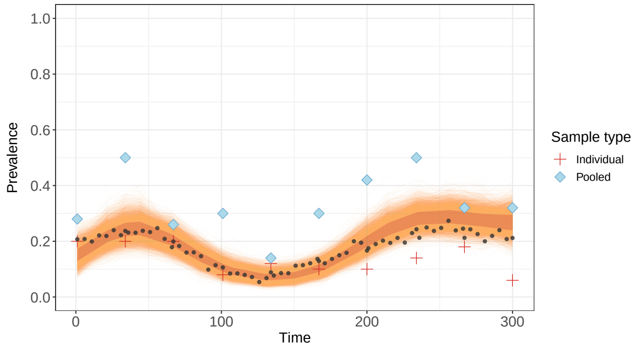

##### False negative rate

False negative rate posterior density with 95% CrI:
True value = 0.1.

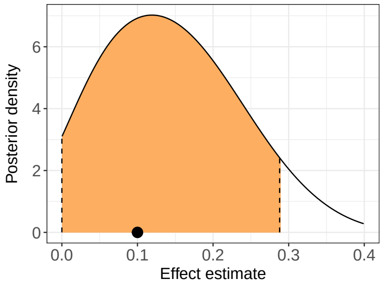

##### Individual covariate

Effect estimate posterior density with 95% CrI, moderately correlating covariate.
True coefficient = 1.9980711.

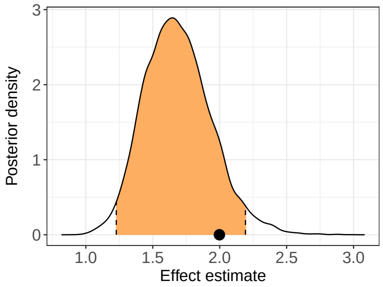

### Irregular sampling times

These datasets have the same prevalence dynamics, sampling sizes and number of sampling sessions as the main dataset, but have different session times and intervals between sessions.
The models use the weakly correlating individual covariate.
There are two versions of this scenario.

#### Irregular sampling times v1

##### Prevalence dynamics

Estimated prevalence dynamics for the main dataset, using a moderately correlating covariate. Black points = simulated prevalence. Orange lines = 5000 posterior draws. Orange band = 50% credible interval.

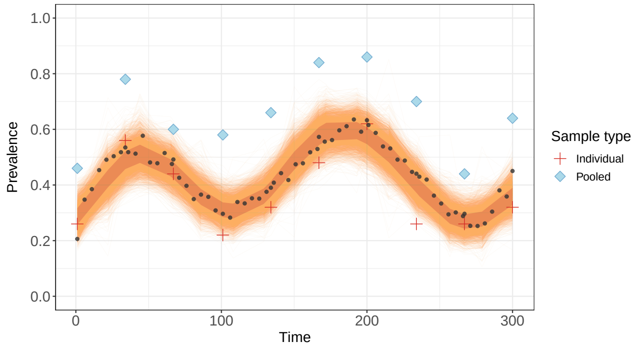

##### False negative rate

False negative rate posterior density with 95% CrI:
True value = 0.1.

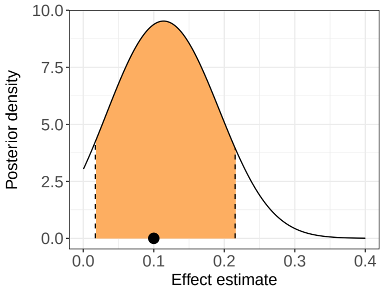

##### Individual covariate

Effect estimate posterior density with 95% CrI, moderately correlating covariate.
True coefficient = 1.8615227.

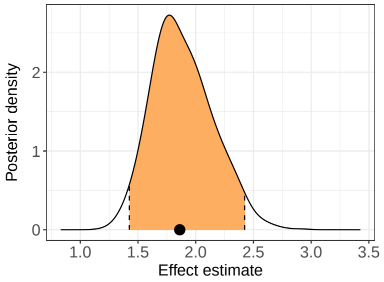
Posterior density
0
1
2
1.0
1.5
2.0
2.5
3.0
3.5
Effect estimate

#### Irregular sampling times v2

With weakly correlating individual covariate.

##### Prevalence dynamics

Estimated prevalence dynamics for the main dataset, using a moderately correlating covariate. Black points = simulated prevalence. Orange lines = 5000 posterior draws. Orange band = 50% credible interval.

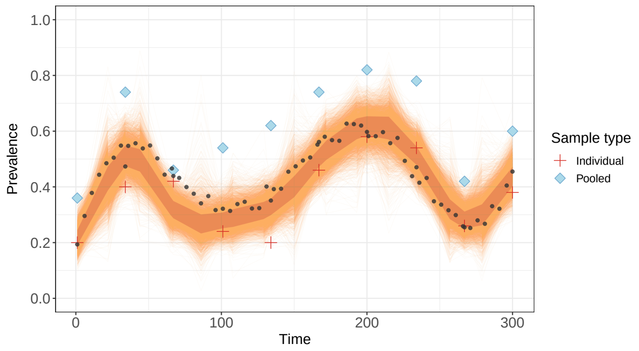

##### False negative rate

False negative rate posterior density with 95% CrI:
True value = 0.1.

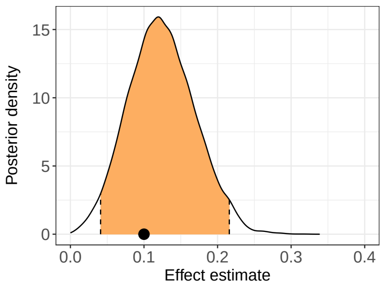

##### Individual covariate

Effect estimate posterior density with 95% CrI, moderately correlating covariate.
True coefficient = 2.2286597.

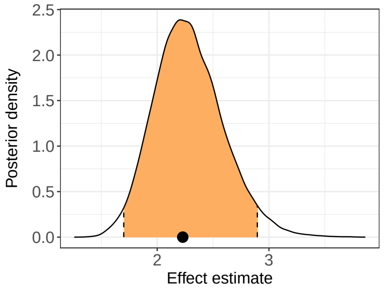

### Lower sample size

These datasets have the same sample timing and number sessions as the main dataset, but the number of samples of each type per sampling session was reduced to 30 (from 50).
The models use the weakly correlating individual covariate.

#### Prevalence dynamics

Estimated prevalence dynamics for the main dataset, using a moderately correlating covariate. Black points = simulated prevalence. Orange lines = 5000 posterior draws. Orange band = 50% credible interval.

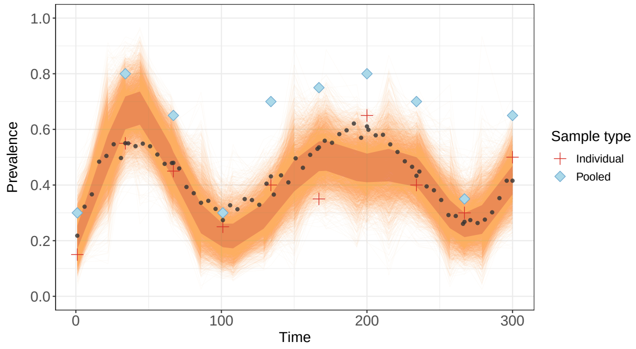

#### False negative rate

False negative rate posterior density with 95% CrI:
True value = 0.1.

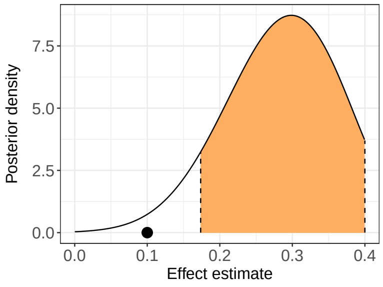

#### Individual covariate

Effect estimate posterior density with 95% CrI, moderately correlating covariate.
True coefficient = 2.9114577.

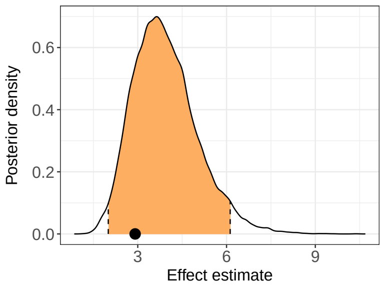

### Fewer sampling sessions

These datasets have the same sample timing and per-session sample size as the main dataset, but the number of sampling sessions was reduced to 5 (from 10).

The models use the weakly correlating individual covariate.

#### Prevalence dynamics

Estimated prevalence dynamics for the main dataset, using a moderately correlating covariate. Black points = simulated prevalence. Orange lines = 5000 posterior draws. Orange band = 50% credible interval.

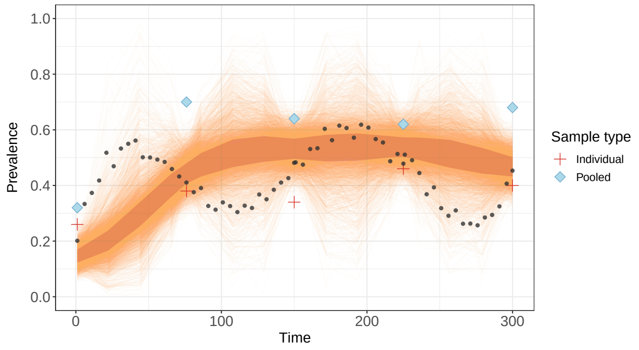

#### False negative rate

False negative rate posterior density with 95% CrI:
True value = 0.1.

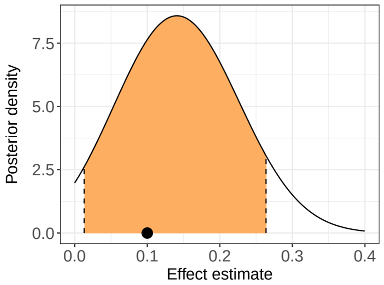

#### Individual covariate

Effect estimate posterior density with 95% CrI, moderately correlating covariate.
True coefficient = 1.519791.

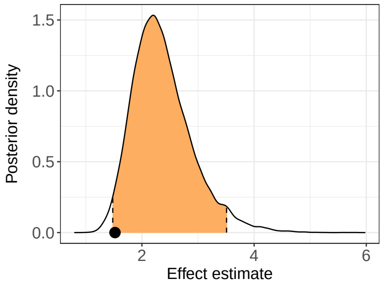

1.5
1.0
0.5
0.0
Posterior density
2
4
6
Effect estimate

### Misspecified Ct distribution

These datasets are the same as the main dataset, except that the Ct values of the pooled samples were simulated assuming a distribution of Ct values in the population that is different from the one used for calculating the probability of observing a certain Ct value (which assumes a low-to-high Ct distribution). This is done for two scenarios: (1) simulations done using a flat distribution and (2) simulations done using a high-to-low distribution.
The models use the weakly correlating individual covariate.

#### Misspecified Ct distribution v1: flat

Ct frequency distribution used to simulate pooled Ct values:

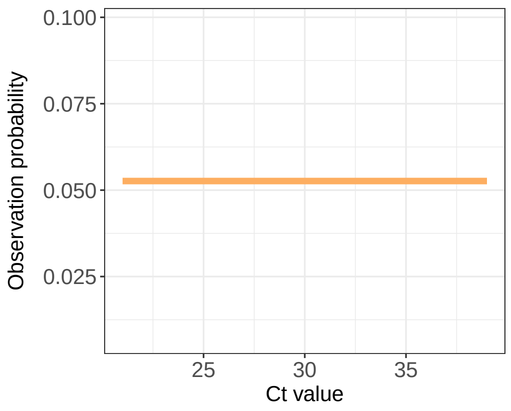

##### Prevalence dynamics

Estimated prevalence dynamics for the main dataset, using a moderately correlating covariate. Black points = simulated prevalence. Orange lines = 5000 posterior draws. Orange band = 50% credible interval.

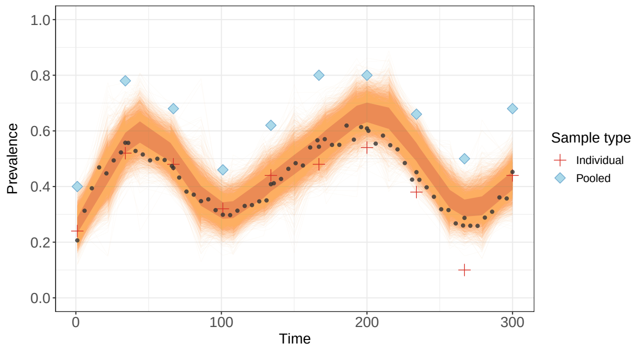

##### False negative rate

False negative rate posterior density with 95% CrI:
True value = 0.1.

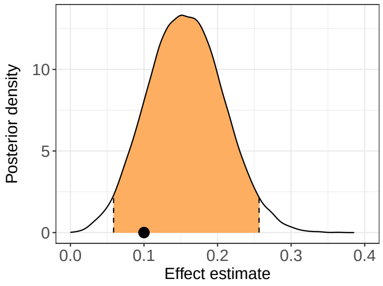

##### Individual covariate

Effect estimate posterior density with 95% CrI, moderately correlating covariate.
True coefficient = 2.3995469.

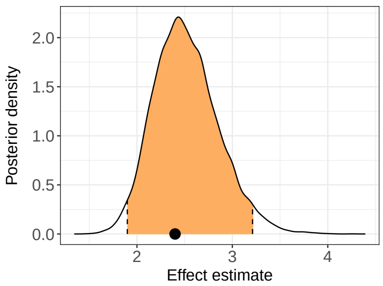

#### Misspecified Ct distribution v2: high-to-low

Ct frequency distribution used to simulate pooled Ct values:

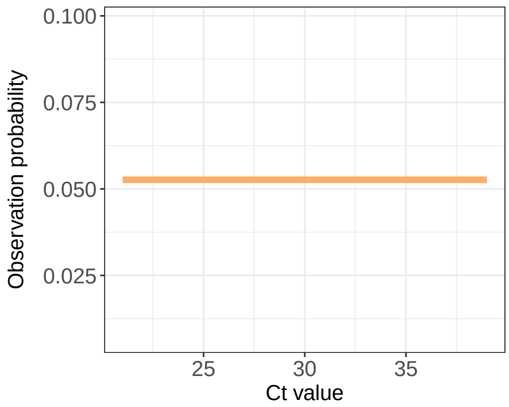

##### Prevalence dynamics

Estimated prevalence dynamics for the main dataset, using a moderately correlating covariate. Black points = simulated prevalence. Orange lines = 5000 posterior draws. Orange band = 50% credible interval.

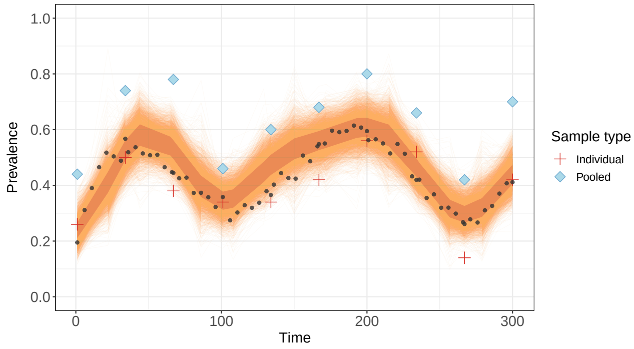

##### False negative rate

False negative rate posterior density with 95% CrI:
True value = 0.1.

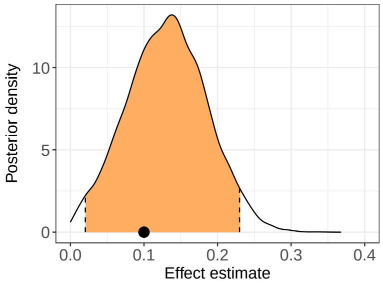

##### Individual covariate

Effect estimate posterior density with 95% CrI, moderately correlating covariate.
True coefficient = 2.1992324.

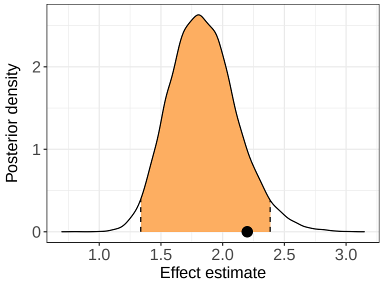

### Misspecified bat count

These datasets are the same as the main dataset.
The model however receives bat count data (the number of bats contributing to a pooled sample) with errors.
This is done for varying degrees of errors (small to large).

The model uses the weakly correlating individual covariate.

#### Misspecified bat count v1

For each count, there is a 50% probability of wrongly decreasing or increasing the bat count "observation".
There is a 30% probability of a 1-count error, and a 20% probability of a 2-count.
If the generated error would result in a count $< 1$, the error is added instead of subtracted.

##### Prevalence dynamics

Estimated prevalence dynamics for the main dataset, using a moderately correlating covariate. Black points = simulated prevalence. Orange lines = 5000 posterior draws. Orange band = 50% credible interval.

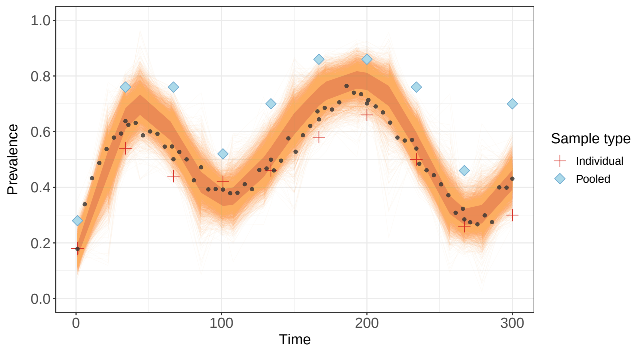

##### False negative rate

False negative rate posterior density with 95% CrI:
True value = 0.1.

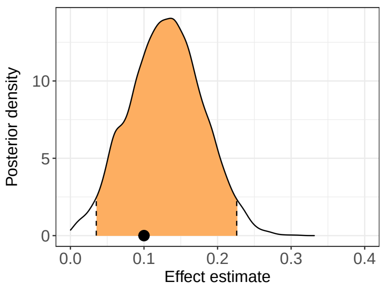

##### Individual covariate

Effect estimate posterior density with 95% CrI, moderately correlating covariate.
True coefficient = 1.7966601.

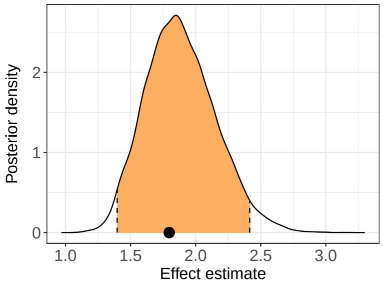
Posterior density
0
1
2
1.0
1.5
2.0
2.5
3.0
Effect estimate

#### Misspecified bat count v2

For each count, there is a 80% probability of wrongly decreasing or increasing the bat count "observation". There is a 50% probability of a 1-count error, and a 30% probability of a 2-count.
If the generated error would result in a count $< 1$, the error is added instead of subtracted.

##### Prevalence dynamics

Estimated prevalence dynamics for the main dataset, using a moderately correlating covariate. Black points = simulated prevalence. Orange lines = 5000 posterior draws. Orange band = 50% credible interval.

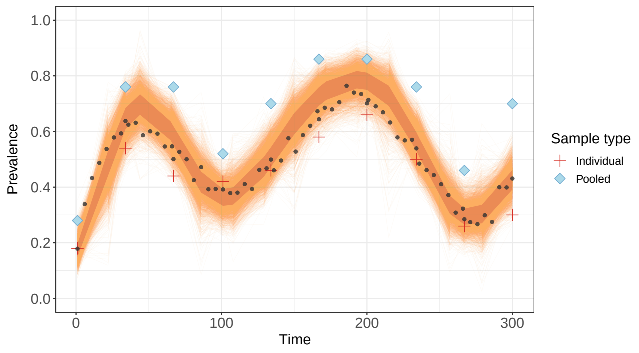

##### False negative rate

False negative rate posterior density with 95% CrI:
True value = 0.1.

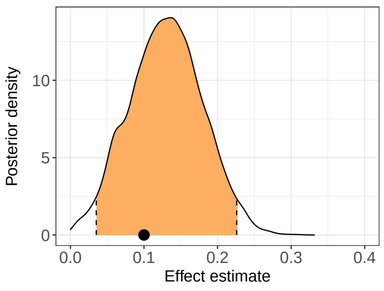

##### Individual covariate

Effect estimate posterior density with 95% CrI, moderately correlating covariate.
True coefficient = 1.7966601.

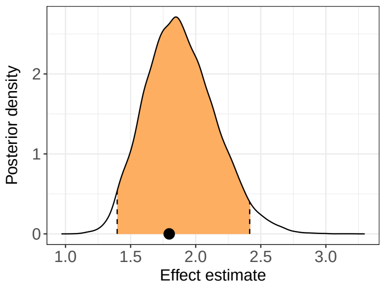

#### Misspecified bat count v3

For each count, there is a 80% probability of wrongly decreasing or increasing the bat count "observation". There is a 20% probability of a 2-count error, 20% 3-count, 20% 4-count, 20% 5-count If the generated error would result in a count < 1, the error is added instead of subtracted.

##### Prevalence dynamics

Estimated prevalence dynamics for the main dataset, using a moderately correlating covariate. Black points = simulated prevalence. Orange lines = 5000 posterior draws. Orange band = 50% credible interval.

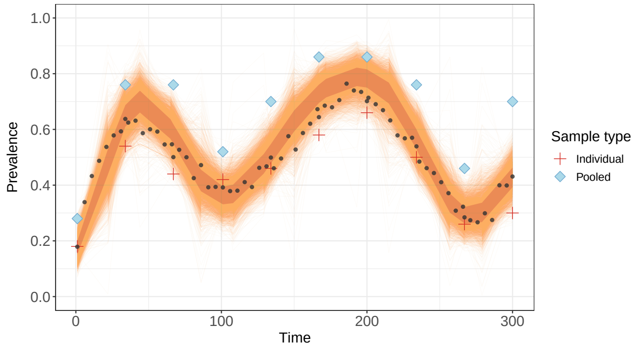

##### False negative rate

False negative rate posterior density with 95% CrI:
True value = 0.1.

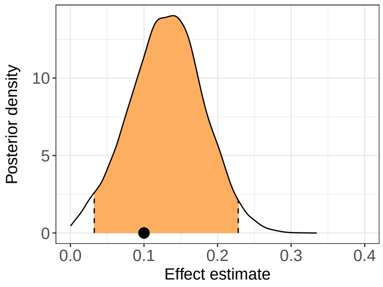

##### Individual covariate

Effect estimate posterior density with 95% CrI, moderately correlating covariate.
True coefficient = 1.7966601.

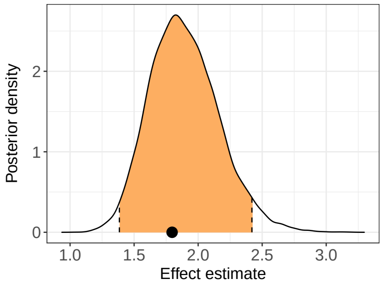

### Prevalence regression

This dataset is the same as the main dataset, with the addition of a covariate that correlates with population-level prevalence, and is estimate using Gaussian Process regression.

The model uses the weakly correlating individual covariate.

Prevalence covariate values for each sampling session were simulated by randomly drawing a value from a normal distribution with a mean equal to the standard cumulative distribution function of prevalence divided by 3.4 (an arbitrarily chosen coefficient), and standard deviation 0.02.

#### Prevalence dynamics

Estimated prevalence dynamics for the main dataset, using a moderately correlating covariate. Black points = simulated prevalence. Orange lines = 5000 posterior draws. Orange band = 50% credible interval.

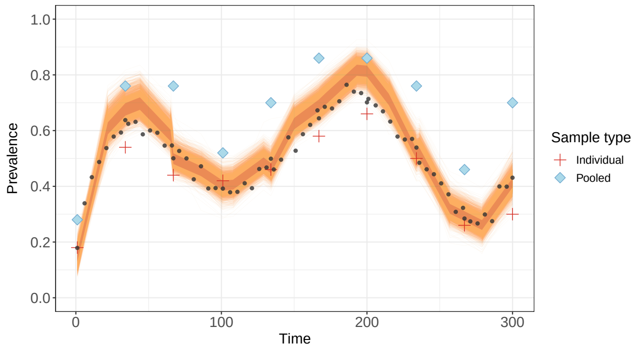

#### False negative rate

False negative rate posterior density with 95% CrI:
True value = 0.1.

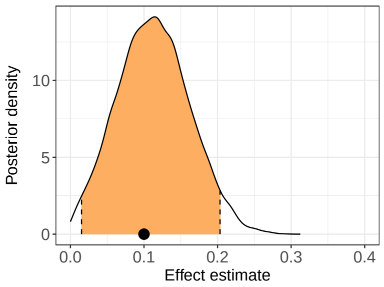

#### Individual covariate

Effect estimate posterior density with 95% CrI, moderately correlating covariate.
True coefficient = 1.7966601.

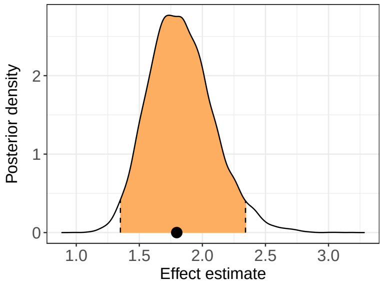

##### Prevalence covariate

Effect estimate posterior density with 95% CrI.
True coefficient = 3.4.

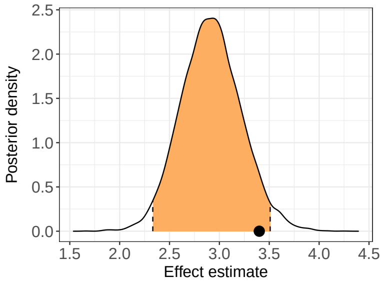

### Asynchronous pooled and individual samples

This dataset has the same prevalence dynamics, sampling sizes and number of sampling sessions as the main dataset, but the timing of pooled and individual sampling is alternated.
The models use the weakly correlating individual covariate.

#### Prevalence dynamics

Estimated prevalence dynamics for the main dataset, using a moderately correlating covariate. Black points = simulated prevalence. Orange lines = 5000 posterior draws. Orange band = 50% credible interval.

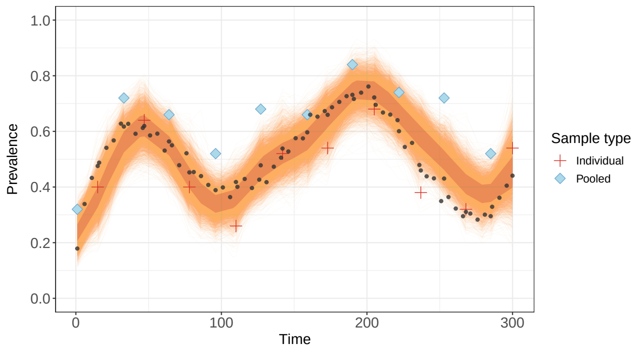

#### False negative rate

False negative rate posterior density with 95% CrI:
True value = 0.1.

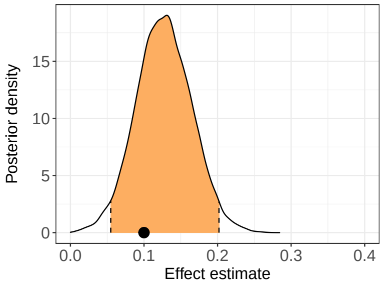

#### Individual covariate

Effect estimate posterior density with 95% CrI, moderately correlating covariate.
True coefficient = 2.276739.

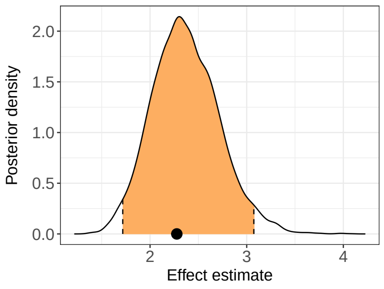

### Irregular and asynchronous sampling times, lower sample sizes (20)

This dataset has the same prevalence dynamics and number of sampling sessions as the main dataset, but the timing of pooled and individual sampling is irregular and asynchronous, and sampling sizes per session are lower (20).
The models use the weakly correlating individual covariate.

#### Prevalence dynamics

Estimated prevalence dynamics for the main dataset, using a moderately correlating covariate. Black points = simulated prevalence. Orange lines = 5000 posterior draws. Orange band = 50% credible interval.

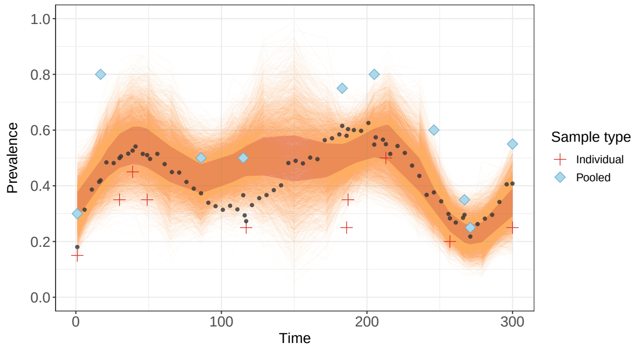

#### False negative rate

False negative rate posterior density with 95% CrI:
True value = 0.1.

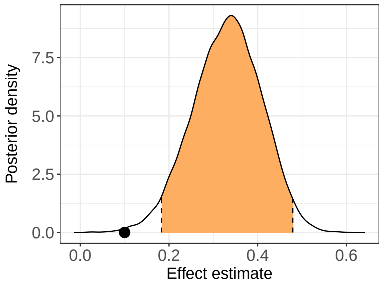

#### Individual covariate

Effect estimate posterior density with 95% CrI, moderately correlating covariate.
True coefficient = 2.1572309.

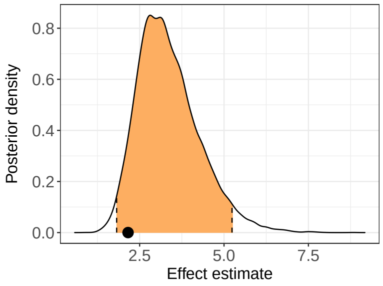
0.8
0.6
0.4
0.2
0.0
Posterior density
2.5
5.0
7.5
Effect estimate

#### Irregular and asynchronous sampling times, fewer sessions

This dataset has the same prevalence dynamics and sample sizes per session as the main dataset, but the timing of pooled and individual sampling is irregular and asynchronous, and there are fewer sampling sessions (5).
The models use the weakly correlating individual covariate.

##### Prevalence dynamics

Estimated prevalence dynamics for the main dataset, using a moderately correlating covariate. Black points = simulated prevalence. Orange lines = 5000 posterior draws. Orange band = 50% credible interval.

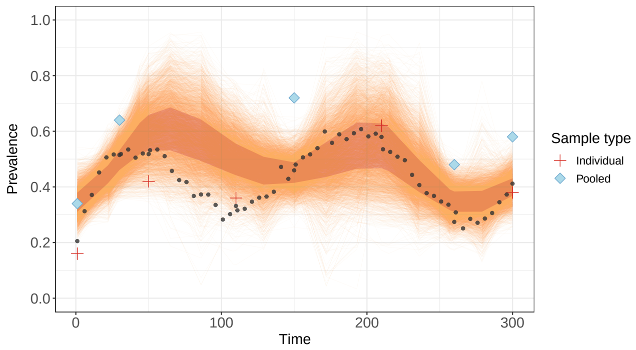

##### False negative rate

False negative rate posterior density with 95% CrI:
True value = 0.1.

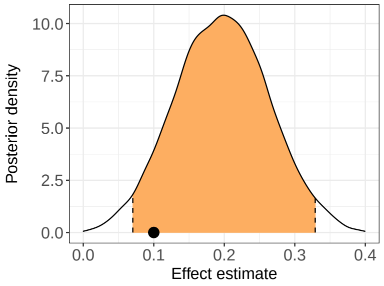

##### Individual covariate

Effect estimate posterior density with 95% CrI, moderately correlating covariate.
True coefficient = 1.7853004.

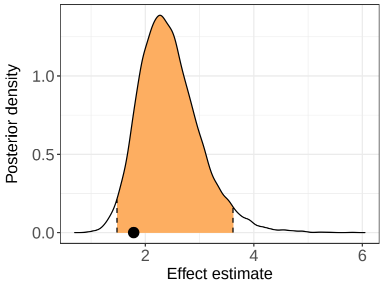

Posterior density
1.0
0.5
0.0
2
4
6
Effect estimate

### Additional figures

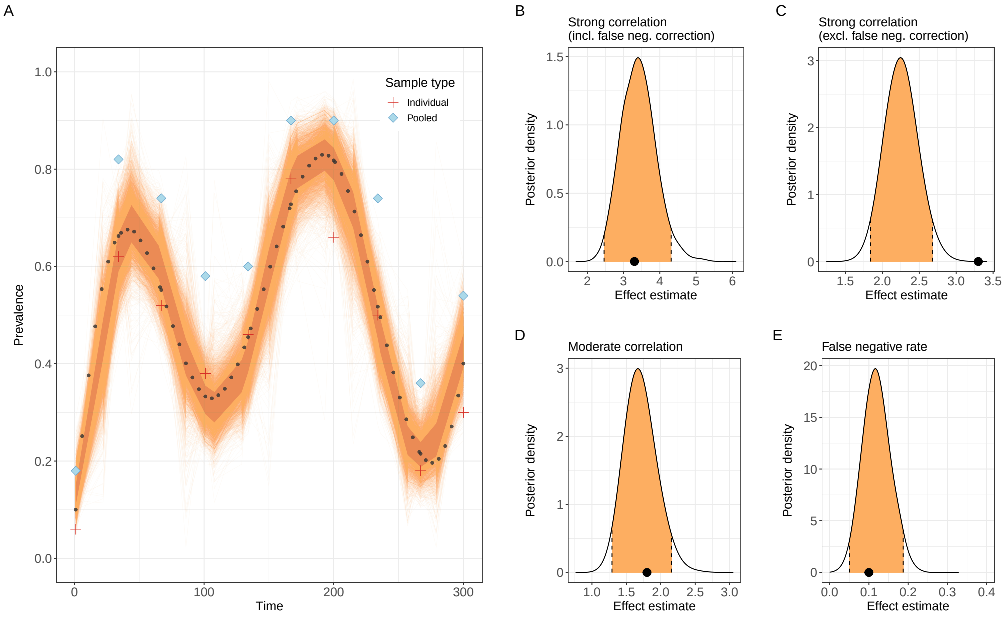

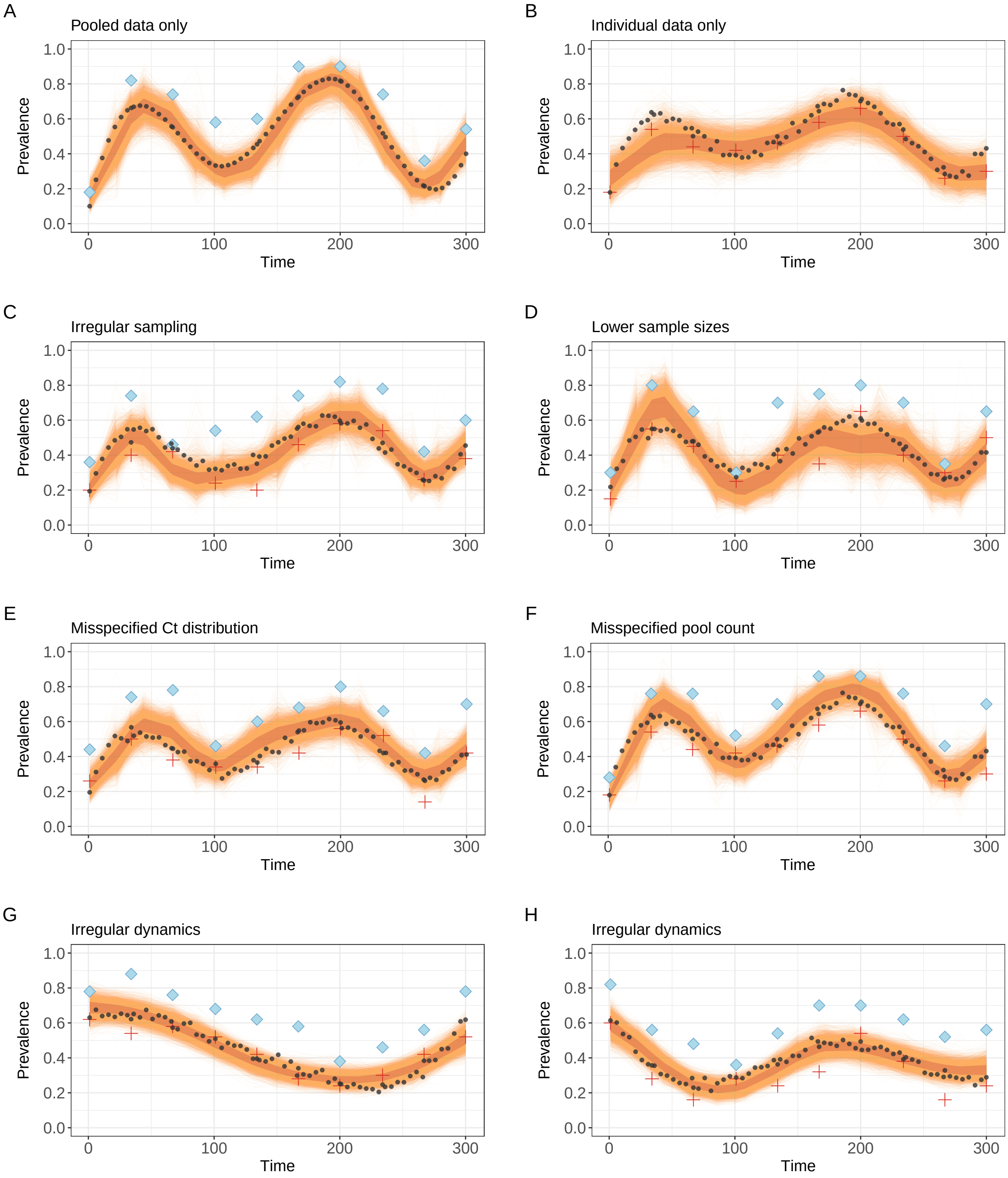
A
Pooled data only
B
Individual data only
C
Irregular sampling
D
Lower sample sizes
E
Misspecified Ct distribution
F
Misspecified pool count
G
Irregular dynamics
H
Irregular dynamics
Prevalence
Time
0.0
0.2
0.4
0.6
0.8
1.0
0
100
200
300
